# Intercellular signaling dynamics from a single cell atlas of the biomaterials response

**DOI:** 10.1101/2020.07.24.218537

**Authors:** Christopher Cherry, David R Maestas, Jin Han, James I Andorko, Patrick Cahan, Elana J Fertig, Lana X Garmire, Jennifer H Elisseeff

**Affiliations:** Translational Tissue Engineering Center, Wilmer Eye Institute and the Department of Biomedical Engineering, Johns Hopkins University School of Medicine, Baltimore, MD; Department of Biomedical Engineering and Institute for Cell Engineering, Johns Hopkins University School of Medicine, Baltimore, MD, USA; Department of Oncology, Johns Hopkins University School of Medicine, Baltimore, MD, USA; Department of Applied Mathematics and Statistics, Johns Hopkins University, Baltimore, MD, USA; Department of Computational Medicine and Bioinformatics, University of Michigan, Ann Arbor. MI 48105; Bloomberg~Kimmel Institute for Cancer Immunotherapy and Sidney Kimmel Comprehensive Cancer Center, Johns Hopkins University School of Medicine, Baltimore, MD

## Abstract

Biomaterials serve as the basis of implants, tissue engineering scaffolds, and multiple other biomedical therapeutics. New technologies, such as single cell RNA sequencing (scRNAseq), are enabling characterization of the biomaterial response to an unprecedented level of detail, facilitating new discoveries in the complex cellular environment surrounding materials. We performed scRNAseq and integrated data sets from multiple experiments to create a single cell atlas of the biomaterials response that contains 42,156 cells from biological extracellular matrix (ECM)-derived and synthetic polyester (polycaprolactone, PCL) scaffold biomaterials implanted in murine muscle wounds. We identified 18 clusters of cells, including natural killer (NK) cells, multiple subsets of fibroblasts, and myeloid cells, many of which were previously unknown in the biomaterial response. To determine intra and intercellular signaling occurring between the numerous cell subsets, including immune-stromal interactions in the biomaterial response, we developed Domino (github.com/chris-cherry/domino), a computational tool which allows for identification of condition specific intercellular signaling patterns connected to transcription factor activation from single cell data. The Domino networks self-assembled into signaling modules and cellular subsets involved in signaling independent of clustering, defining interactions between immune, fibroblast, and tissue-specific modules with biomaterials-specific communication patterns. Further compilation and integration of biomaterials single cell data sets will delineate the impact of materials chemical and physical properties and biological factors, such as anatomical placement, age, or systemic disease, that will direct biomaterials design.

## Introduction

Biomaterials serve as the basis of medical devices that first entered into clinical practice in the 1960s. After evidence of the body’s reaction to biomaterials emerged, the foreign body response (FBR) was defined and characterized. The FBR process initiates with an immune response and may develop into chronic inflammation and fibrosis around the implant (Anderson, 2001; Anderson et al., 2008). This fibrosis is a major challenge in the design of therapeutic biomaterials. As the field of regenerative medicine and tissue engineering developed decades later, biomaterial design turned to building scaffolds that created a desired biological response, such as mobilizing stem cells, promoting vascularization and stimulating tissue development instead of causing FBR (Zakrzewski et al., 2014). Biomaterial scaffolds are also employed as a tool to engineer tissue models to probe mechanisms of tissue development and disease, such as cancer in controlled three dimensional environments (Zhang et al., 2018). While previous foundational research described a number of key cell responses to biomaterials, recent technology development now enables an unprecedented ability to map cell responses and tissue composition. Armed with a comprehensive understanding of the cells responding to biomaterials, the local tissue environment, and cell interactions, we can better design biomedical implants and tissue engineering scaffolds to avoid FBR and understand mechanisms of clinical adverse events.

Developments in high parametric flow cytometry are expanding the tools available to evaluate which cells are responding to implantation of biomaterials based on expression of protein surface markers characteristic of a specific cell type. This technology enabled the discovery of more rare cell types responding to biomaterial implants, such as, CD4+ T helper cells expressing interleukin (IL)-4 and IL17 and B cells (Chung et al., 2020; Sadtler et al., 2016). Single cell RNA sequencing (scRNAseq) is another method that can be applied to evaluate biomaterial responses that uses unbiased identification of groups of cells based on gene expression without any *a priori* knowledge of cellular phenotypes or markers (Papalexi and Satija, 2018). The resulting clusters of cells can be phenotyped based on their gene expression profiles and mapped back to experimental condition. scRNAseq has transformed experimental approaches and understanding in cancer (Suvà and Tirosh, 2019), autoimmunity (Das et al., 2018; Zhang et al., 2019), and infection responses (Steuerman et al., 2018; Yao et al., 2019) but has not been widely applied to biomaterials. Application of scRNAseq to the biomaterials response will provide new understanding of cellular response that may transform their design.

Cells from the innate and adaptive immune system respond to biomaterials when they are implanted in the body. Subsequently, fibroblasts are activated and programs of fibrosis and/or tissue repair develop. The communication network between the immune and stromal compartments in the biomaterial response is unknown. The application of scRNAseq to the biomaterials response introduces even more cell types that contribute to the biomaterial response, creating a complex network of communication in the tissue. Reconstruction of cellular communication networks (ligand-receptor activity) from scRNAseq data sets can provide insight into tissue microenvironments and physiological responses despite missing cell types and limited sequencing depth to elucidate biological mechanisms and potential therapeutic targets. Computational tools currently available for ligand-receptor signaling analysis of scRNAseq data sets correlate expression of ligands and receptors across groups of cells with no phenotypic connection (Efremova et al., 2020; Noël et al., 2020; Wang et al., 2019) or require an *a priori* gene set (Browaeys et al., 2020). Use of this method to identify signaling pathways connected with downstream phenotypic changes is not possible without a previously defined gene set which is not currently available for biomaterial responses.

Here, we applied scRNAseq to biological and synthetic biomaterial scaffolds implanted in muscle defects. We used integration techniques similar to those from the Human Cell Atlas (Regev et al., 2017) and the Tabula Muris Consortium (Schaum et al., 2018) to pool multiple single cell data sets including sorted cells and to provide a single cell atlas of response to biomaterials. We believe the atlas and analysis techniques detailed here will provide a foundation for integration with data sets from other biomaterials and tissues in the future. The resulting atlas contains ~42,000 cells from a single murine wound model treated with both biologic and synthetic biomaterials. We identified new cell types responding to synthetic and biological biomaterials and determined cell types not captured for single cell analysis through flow cytometry comparisons. To reconstruct cell communication networks that develop in response to biomaterials, we developed Domino, a computational program that reconstructs intracellular communication based on transcription factor activation. We link transcription factor activation with expression of receptors and their possible ligands. This process generates signaling hypotheses with specific, testable downstream biological function through transcription factor activation in specific cell types. Application of Domino to the biomaterials atlas defined signaling modules for stromal, immune, and tissue-specific cells independent of clustering. The signaling predictions identified validated signaling and identified new pathways associated with regeneration and fibrotic wound healing. To validate Domino’s use in other contexts, we applied it to a previously published dataset from Alzheimer’s patients (Grubman et al., 2019) and identified new signaling pathways specific to diseased or healthy brain.

## Results

### A single cell atlas of the biomaterials tissue microenvironment

To construct a single cell atlas of the biomaterials microenvironment, we integrated single cell RNA sequencing (scRNAseq) data sets from multiple experiments and different cell preparations. Synthetic and biological scaffolds were implanted in a murine model of volumetric muscle loss (VML) and compared to wound alone. The VML injury provides a combination of tissue damage and biomaterial response and provides a pocket for biomaterial implantation (Sicari et al., 2012). A biological scaffold derived from the extracellular matrix (ECM) of small intestinal submucosa (SIS) served as a model for a scaffold used in preclinical and clinical application of tissue repair that induces a Type 2 immune environment (Badylak and Gilbert, 2008; Sadtler et al., 2016). We evaluated polycaprolactone (PCL) as a model synthetic biomaterial that is used preclinically and clinically in devices. PCL induces a Type 17 immune environment and fibrosis (Anderson et al., 2008; Chung et al., 2020). When the ECM and PCL materials are implanted in muscle, they induce significant changes in gene expression in the bulk tissue as measured by Nanostring™ after 1 week (Fig. 1A). Of the 770 genes that were tested, 397 were differentially expressed between PCL and saline and 194 between ECM and saline. Of the differentially expressed genes, 216 were unique to PCL and 13 were unique to ECM, confirming that the biomaterials induced different responses at the global tissue level (Supplementary Table 1).

**Figure 1:**
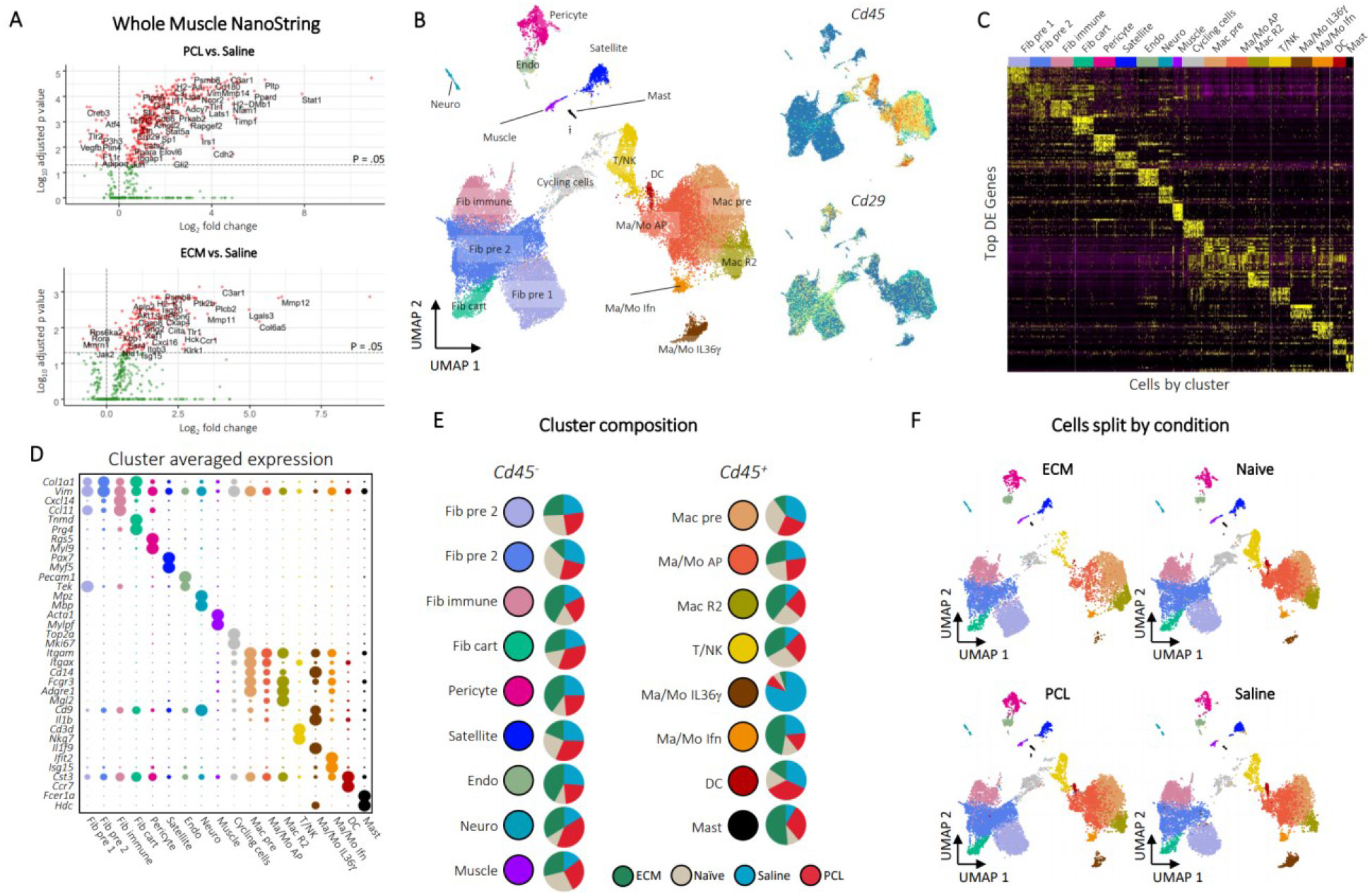
A Single Cell Atlas of the Biomaterials Immune Microenvironment. **A**, NanoString gene expression analysis of mRNA isolated from whole muscle samples of animals treated ECM or PCL compared to saline treated animals. nSolver© software with default parameters was used for statistical testing with adjusted *P* ≤ 0.05 defining differential expression. **B**, Overview of cell clusters identified in the composite scRNAseq dataset. UMAP plots with cluster labels (left) and gene expression levels of *Cd45* and *Cd29* (right) are shown. **C**, Heatmap of up to 10 differentially expressed genes with highest log fold-change from each cluster. Cells are ordered and labeled by cluster with random sampling of up to 100 cells per cluster to ensure visibility of small clusters. **D**, Gene markers for single cell subsets. The dotplot shows expression of genes associated with cluster identity. Cluster averaged gene expression values after normalization to the maximum averaged expression are shown. **E**, Composition of cluster by condition. Cluster labels and colors are shown adjacent to pie charts of normalized number of cells by condition. Prior to comparison of conditions, cells were normalized to the total number CD45^+^ or CD45^-^ cells by sample. The resulting values are compared across condition. Cycling cells are not shown. Only samples from non-sorted samples were used to calculate proportions (see methods). **F**, Visualization of cells by condition on UMAP. Each UMAP shows cells only from a given condition colored by cluster.

To generate the scRNAseq data sets we created single cell suspensions from the muscle tissue with and without biomaterials for application to 10X and DropSeq. Single cell suspensions from whole tissue preparations were enriched for CD45^+^ cells to enable capture of less frequent cell populations. Fluorescence activated cell sorting (FACS) was used to apply specific cell populations to 10X including macrophages (CD45^+^F4/80^hi+^Ly6c^+^CD64^+^) (Sommerfeld et al., 2019) and mesenchymal/fibroblasts (CD45^-^CD19^-^CD31^-^CD29^+^) (Supplementary Fig. 1A-C). A detailed description of age, time of harvest, treatment, and sorting methodology for each of the samples in the atlas is provided in Supplementary Table 2. The resulting dataset includes 42,156 cells with an average of 198,000 reads and 1,167 genes per cell after filtering low-quality cells and genes. To determine whether the integration of sorted cell data sets caused biases with clustering due to increased numbers of the sorted cell populations, we compared outcomes of the whole tissue analysis with and without the sorted cell data sets (Supplementary Fig. 1D). While inclusion of the macrophage and fibroblast sort data increased these populations and their resolution, cell populations present only in the unsorted dataset maintain clear, distinct phenotypes by both clustering and UMAP. The integrated dataset, including cells from sorted and CD45^+^ enriched whole tissue single cell suspensions, is used for all subsequent analyses.

### Identification of immune and stromal single cell clusters in the biomaterial response

Unsupervised clustering identified 18 distinct clusters: nine CD45^-^ clusters, eight CD45^+^ clusters, and one cluster of cycling cells with both CD45^+^ and CD45^-^ cells (Fig. 1B). We used expression of canonical marker genes, as well as similarity of gene expression profiles, to classify clusters (Fig. 1C, D, Supplementary Table 3). The CD45^+^ clusters included a mixed cluster of T and natural killer (NK) cells (*Cd3d, Ngk7*) (T/NK), dendritic cells (*Ccr7, Cst3*) (DC), and mast cells (*Cpa3, Fcer1a, Hdc*) (Mast), as well as five clusters of monocytes and/or macrophages (*Cd11b, Cd14, Adgre1*) (Fig. 2A-B). The myeloid clusters were composed of an anti-inflammatory macrophage cluster (*Cd163, Retnla*) (Mac R2), a cluster performing antigen presentation (*Cd74, H2-Ab1*) (Mac/Mo AP), a type 17 inflammatory cluster (*Il36γ*) (Mac/Mo Il36γ) (Sommerfeld et al., 2019), and a cluster of cells showing response to interferons (*Ifit2, Isg15*) (Mac/Mo Ifn) (Fig. 2D). The T cell/NK cell subset included cells expressing *Cd4, Cd8, Foxp3*, and *γδ* markers, but there was not adequate resolution to separate these cells into different clusters or differentiate respective subsets (Supplementary Fig. 2). From flow cytometry data, including intracellular staining, there were CD4+ T cells and gamma deltas expressing different cytokines in the biomaterial response that were not captured with scRNAseq (Chung et al., 2020; Sadtler et al., 2016). Further enrichment of these cell types using sorting similar to the macrophage and fibroblast before 10X may be required to capture physiologically relevant resolution of T cells.

**Figure 2:**
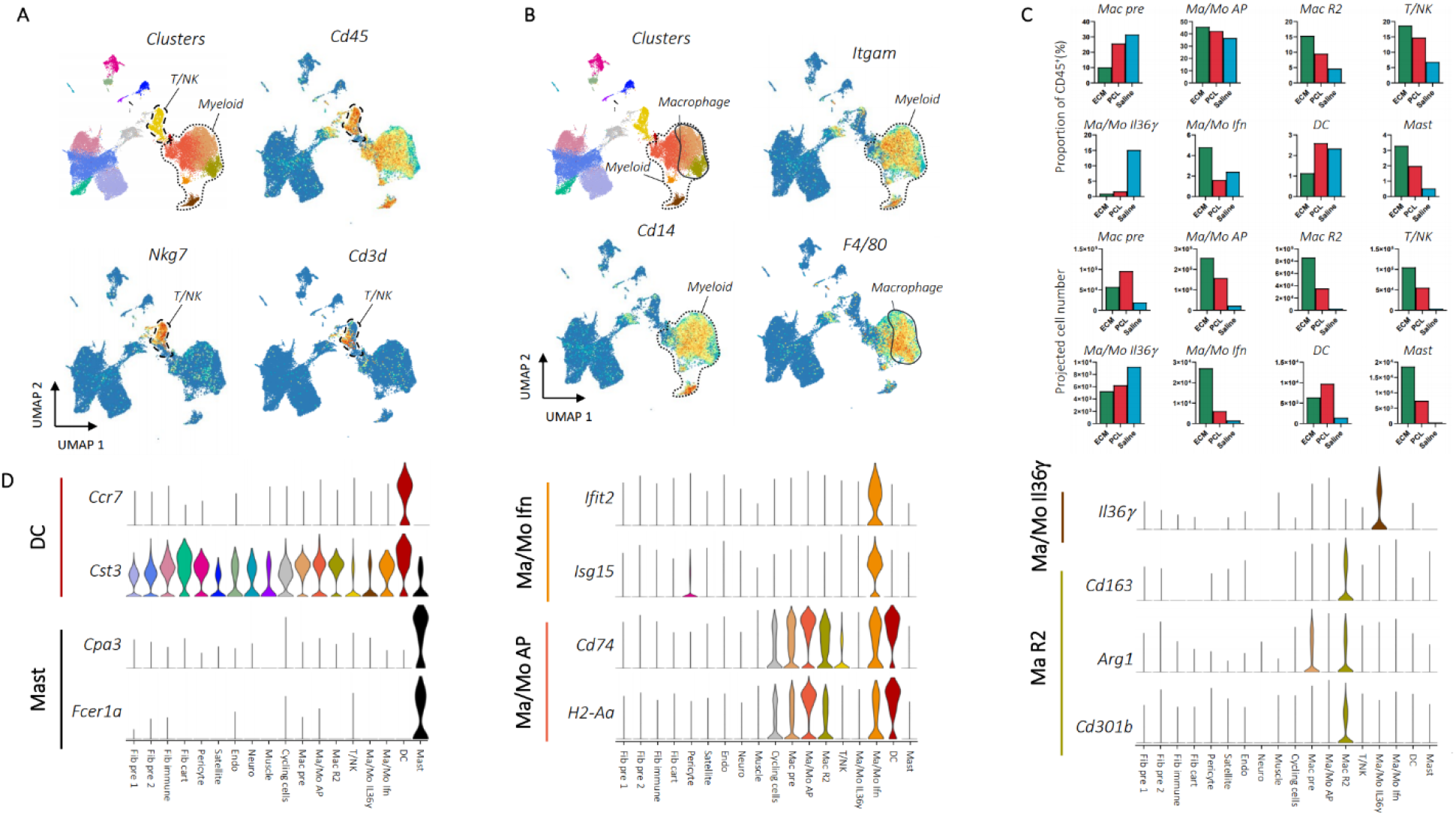
Cd45^+^ scRNAseq cluster expression characteristics. **A**, Immune markers for T and NK cells. Clusters are shown adjacent to *Cd45, Nkg7*, and *Cd3d* gene expression. The T/NK cell cluster and myeloid clusters are encircled with dotted outlines. **B**, Gene expression for myeloid and macrophage cell markers. Clusters are shown adjacent to *Itgam, Cd14*, and *F4/80* gene expression. Myeloid and macrophage clusters are encircled and labeled. **C**, Cluster condition proportions and projection using flow cytometry cell numbers. For each CD45^+^ cluster, the percent of total CD45^+^ cells by condition was calculated (top). The resulting proportions were multiplied by total CD45^+^ cell numbers from flow cytometry in ECM, PCL, or saline treated animals (Supplementary Fig. 3) to obtain projected cell numbers from the scRNAseq proportions (bottom). **D**, Gene expression for myeloid cluster markers. Single cell gene expression for marker genes used to identify myeloid clusters in tandem with CD14, CD11b, and F4/80 expression are shown as violin plots.

The CD45^-^ clustering identified multiple types of mesenchymal, stromal, and fibroblast-like cells. Five clusters were identified as involved in specialized tissue formation, including Schwann cells (*Mpz, Mbp, Plp1*) (Neuro), pericytes (*Rgs5, Acta2, Mcam*) (Pericyte), endothelial cells (*Cd31, Cavin2, Ptprb*) (Endo), satellite cells (*Pax7, Des, Myf5*) (Satellite), and myoblasts (*Acta1, Tnnt3, Mylpf*) (Muscle), as well as four clusters of fibroblasts (*Col1a1, Pdgfra, Postn*) (Fig. 3A). Two of the fibroblast clusters expressed *Osr1* (Fig. 3B), a transcription factor indicating stemness (Vallecillo-García et al., 2017). CytoTRACE, an algorithm used to score cells for stemness, gave high stemness scores to the same two clusters (Fig. 3B), suggesting these two populations serve a stem-like role in the wound environment (Fib pre 1, Fib pre 2). Of the remaining two fibroblast clusters, one appeared cartilage-like based on expression of *Tnmd, Thbs4*, and *Prg4* (Fib cart) and one appeared to be involved in immune regulation based on expression of cytokines *Cxcl14* and *Ccl11* (Fib immune)(Fig. 3C).

**Figure 3:**
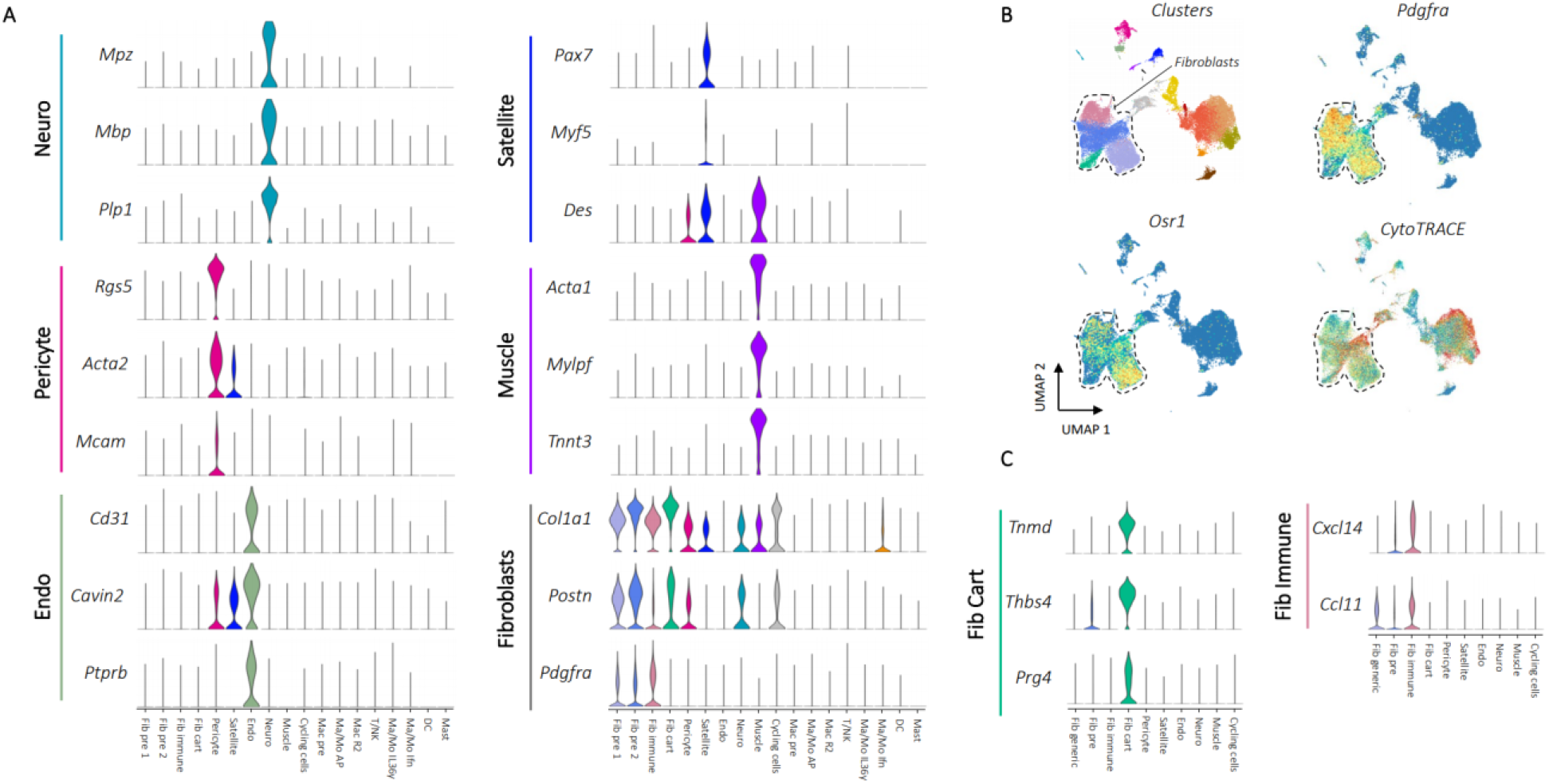
Phenotyping of CD45^-^ scRNAseq clusters. **A**, Marker gene expression for non-fibroblast CD45^-^ cell populations. Up to three characteristic gene markers are shown for each cluster by violin plot of normalized gene expression data. Fibroblast markers were used to identify the four fibroblast clusters Fib pre 1, Fib pre 2, Fib immune, and Fib cart. **B**, Stemness markers for fibroblasts. *Pdgfra* expression and clusters are shown to demonstrate UMAP location of fibroblasts with a dotted line drawn surrounding the fibroblasts. Stem marker *Osr1* and CytoTRACE score, an algorithm used to score cells for stemness, is shown below. **C**, Expression of characteristic markers for the cartilage producing and immune fibroblast clusters.

While single cell identified new populations in the biomaterial response, comparison of the scRNAseq dataset with multiparametric flow cytometry revealed several populations that were missing or underrepresented. Both eosinophils and neutrophils that comprise a large portion of the CD45^+^ population in ECM and PCL, respectively (Supplementary Fig. 3), were not captured in the single cell analysis. Recent work on granuloctyes, including basophils and neutrophils, demonstrate the challenge in capturing these cells and the requirement for additives to preserve their integrity for single cell (Cohen et al., 2018; Szczerba et al., 2019; Xie et al., 2020). The lymphoid populations in our single cell dataset were also limited compared to flow cytometry.

We next examined differences in cellular composition of clusters in the ECM and PCL environments. The number of cells in a particular cluster varied depending on experimental condition and biomaterial (Fig. 1E, F, and Supplementary Table 4). In particular, there were significantly more endothelial cells, mast cells, and anti-inflammatory macrophages (Mac R2) in the ECM implants compared to other conditions. The anti-inflammatory population has been associated with regeneration previously (Sommerfeld et al., 2019), and increased endothelial cell populations may suggest increased vascularization which has been associated with regeneration (Joanisse et al., 2017). The saline condition contained most of the *Il36γ* producing myeloid population. IL36γ producing myeloid populations have been previously shown to be dependent in IL17 signaling in wound healing (Sommerfeld et al., 2019). Cell population comparisons in different experimental groups changed significantly when the total number of CD45^+^ cells from flow cytometry is used to predict total cell number for each of the CD45^+^ clusters (Fig. 2C). Many clusters with similar proportions among the conditions, such as the inflammatory myeloid (Mac-Mono Inflam), have drastic differences in predicted cell number due to the ~10 fold increase in CD45^+^ cell number in the ECM environment compared to saline as determined by flow cytometry (Supplementary Fig. 3B).

While there were proportional differences between some clusters in the different biomaterial environments, additional factors beyond cell composition may be responsible for the phenotypic differences observed experimentally between PCL and ECM environments. We hypothesized that differences in signaling between cells or groups of cells in the different biomaterial-tissue environments may be critical for determining the divergent physiological outcomes in these materials. Furthermore, differences in signaling and transcriptional activation may capture changes resulting from epigenetic and other changes that are not detected in standard single cell expression analysis.

### Development of a computational program to generate inter- and intracellular signaling networks from single cell RNA sequencing data

To further analyze the biomaterial single cell dataset, we developed a program to generate inter- and intracellular signaling networks. Intercellular communication can be modeled as the result of four biological steps (Fig. 4A). Ligands (L) are produced by signaling cells, ligands bind to and activate receptors (R) on target cells, receptor activation triggers a signaling cascade, and transcription factors (TF) at the end of the cascade initiate transcription of specific genes. These transcriptional changes then lead to phenotypic changes in the target cell population. Domino reconstructs cell communication events in reverse to identify signaling connected with specific changes in transcription factor activity (Fig. 4B).

**Figure 4:**
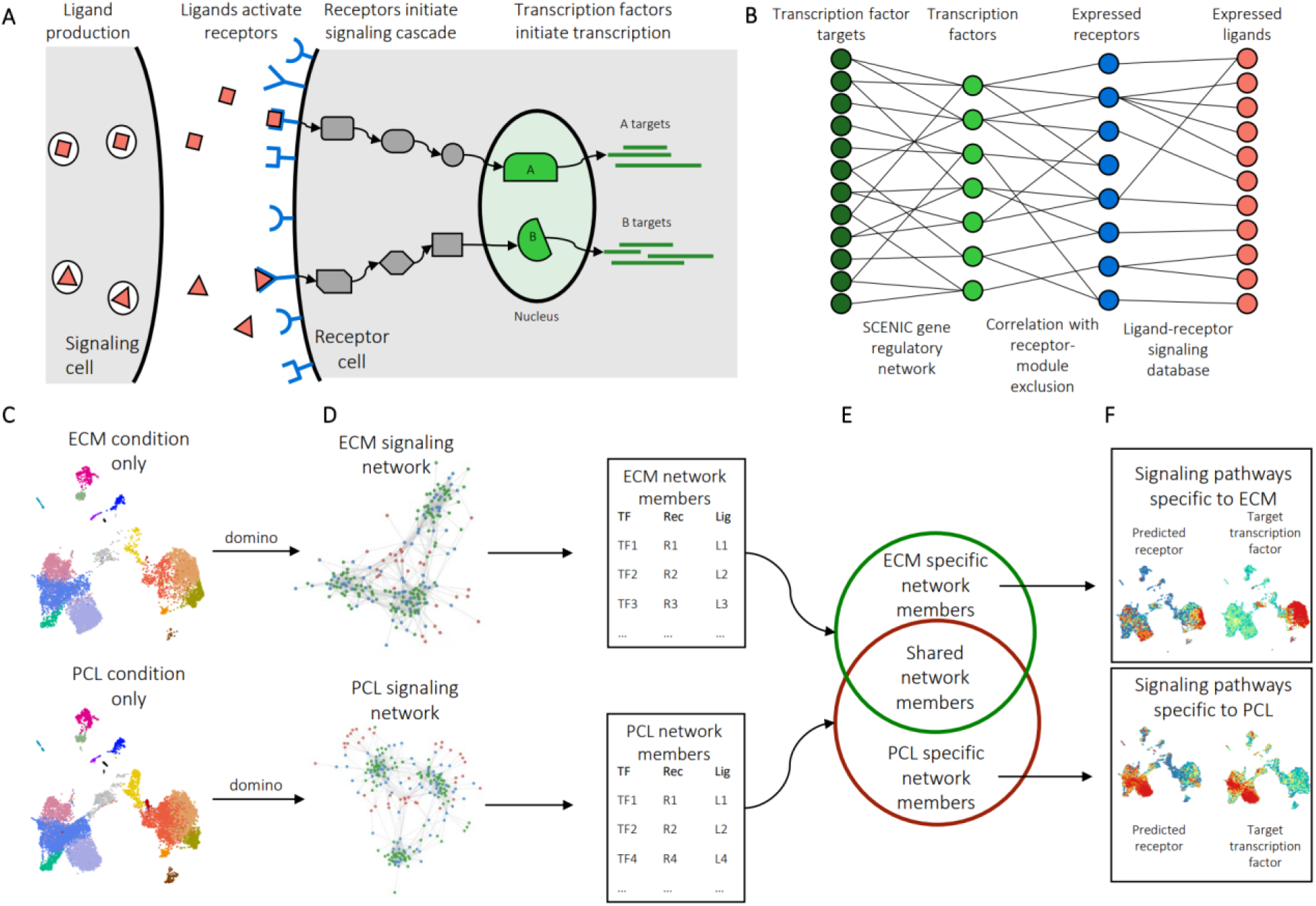
Generation of intercellular and intercluster signaling networks. **A**, A model of biological ligand-receptor signaling. Ligands expressed in a signaling cell bind and activate receptors on a target cell. Subsequent protein-protein signaling triggers activation of transcription factors in the nucleus and expression of target genes. **B**, Reconstruction of a dataset-wide signaling network. SCENIC is used to estimate transcription factor gene regulatory modules, as well as transcription factor activation scores on a cell-by-cell level. Receptors expression levels are correlated with transcription factor activation scores across the entire dataset with exclusion of receptors present in the transcription factor modules. Public receptorligand databases and then queried to identify ligands capable of activation receptors. **C**, Identification of condition specific signaling patterns. In order to identify signaling specific to experimental conditions, the dataset is first split by condition, for example PCL and ECM treated cells. Domino is run on each to identify a signaling network specific to each condition. **D**, Members of signaling networks represent transcription factors, receptors, and ligands that are predicted as active in each condition. **E**, Comparison of network members from each condition identifies transcription factors, receptors, and ligands that are only predicted as active in one condition. **F**, Condition specific members can be used to identify signaling pathways which are only active or are differentially activated in an experimental condition.

The signaling networks that Domino constructs can be used to identify signaling specific to experimental condition. The dataset is first split by condition, for example, cells from ECM-treated tissue and cells from PCL-treated tissue (Fig. 4C). We then generate a signaling network for each treatment, containing all TF, R, and L predicted to be active in each condition (Fig. 4D). We compare the two networks to identify members specific to each (Fig. 4E), identifying TF, R, and L predicted as active in only one condition. The network members active in only one condition can then be used to identify signaling pathways that are specific to that condition (Fig. 4F).

To reconstruct cell communication, Domino first uses the SCENIC gene regulatory network analysis pipeline (Aibar et al., 2017) to generate transcription factor activation scores from raw counts data. SCENIC uses transcription factor expression as input data with a gradient boosting machine (GBM) to predict gene expression of other genes. It extracts co-expression modules from the fitted GBM, prunes the modules for presence of *cis*-regulatory motifs upstream of target genes on the genome, and scores gene regulatory networks on a cell-by-cell basis with area under the recovery curve across the ranking of genes in a cell. Second, Domino connects transcription factor scores with receptor expression using Pearson correlation. Receptors highly correlated with transcription factors are potential candidates for activation of TFs. To ensure correlation is not due to transcription factor targeting of the receptor, Domino excludes receptors found that are predicted as targets of specific transcription factors. Finally, CellphoneDB, a publicly available ligand-receptor database (Efremova et al., 2020), is queried to identify potential ligands for receptors.

The L-R-TF linkages that Domino assembles from the cell communication reconstruction form a global signaling network for the dataset irrespective of any clustering. This independence from clustering allows for unsupervised exploration of L-R-TF activation in single cell data sets. Because Domino starts from activated transcription factors instead of receptors or ligands like other programs, signaling pathways in a target cell population can be identified in the absence of ligand expression which may not be captured by single cell analysis. The process also naturally selects for receptors which are more likely to be activated *in vivo*, rather than looking at all expressed receptors. Further, transcription factors are typically well documented in literature, making connection of specific ligand-receptor pairs with a hypothesized biological change possible. While generation of the signaling network is independent of clustering information, cluster labels can be used to generate an intercluster signaling network if desired (Supplementary Fig. 4)

### Domino identifies signaling patterns associated with biomaterial conditions

We then applied Domino to the biomaterials atlas to determine global signaling networks for the ECM (Fig. 5A) and PCL (Fig. 5D) tissue environments. Labeled, high resolution networks are available as supplementary files. Both the ECM and PCL signaling networks, receptors, and transcription factors self-assembled into modules of signaling in fibroblasts, immune cells, or tissue-specific cells when visualized with a force-directed graph. This self-organization represents unsupervised assembly of both signaling patterns, as well as groups of cells involved in each signaling pathway, and demonstrates the importance of communication between immune-stromal-tissue cell modules.

**Figure 5:**
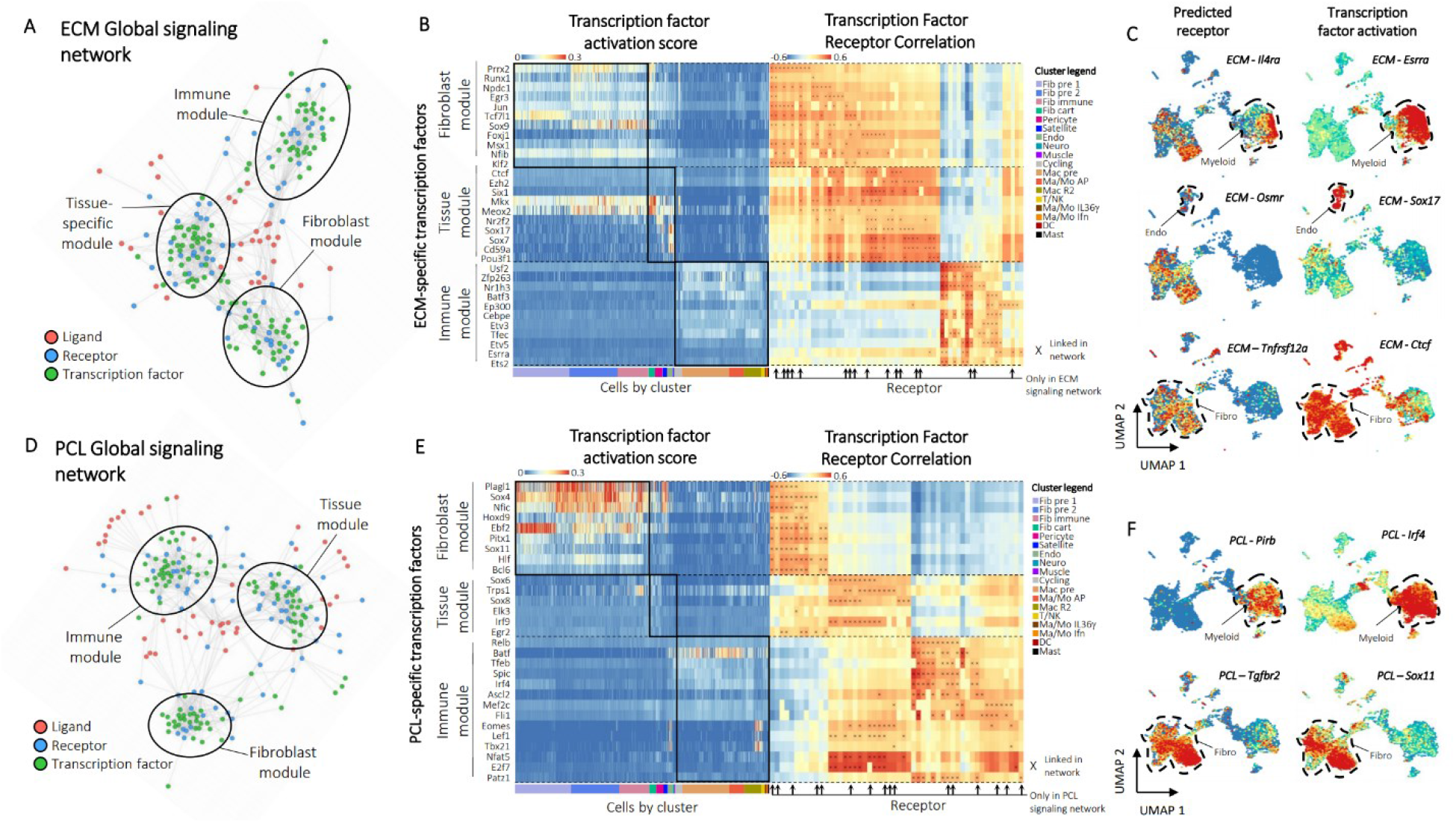
Domino identifies biomaterials condition specific signaling. **A**, The ECM global signaling network. Three modules of receptors and transcription factors are readily apparent and labeled based on enrichment of transcription factors by cluster. **B**, Heatmaps of transcription factor activation score for ECM-specific transcription factors (left) and correlation of transcription factor activation score with receptor expression (right). Transcription factors are binned according to their membership to the fibroblast, tissue, or immune modules from the ECM global signaling network. Cells are ordered and colored according to their cluster. Receptors found only in the ECM condition are marked with arrows. Connections between receptor and transcription factors are marked with an ‘x’ on the correlation heatmap. **C**, Example feature plots of gene expression and activation scores for specific receptor-transcription factor pairs identified by domino in the ECM condition. **D**, The PCL global signaling network. Three modules of receptors and transcription factors are readily apparent and labeled based on enrichment of transcription factors by cluster. **E**, Heatmaps of transcription factor activation score for PCL-specific transcription factors (left) and correlation of transcription factor activation score with receptor expression (right). Transcription factors are binned according to their membership to the fibroblast, tissue, or immune modules from the PCL global signaling network. Cells are ordered and colored according to their cluster. Receptors found only in the PCL condition are marked with arrows. Connections between receptor and transcription factors are marked with an ‘x’ on the correlation heatmap. **F**, Example feature plots of gene expression and activation scores for specific receptor-transcription factor pairs identified by domino in the fibrotic (PCL) condition.

ECM specific transcription factors were enriched in fibroblasts, tissue-specific cells, and immune cells according to their modules as predicted by Domino (Fig. 5B). Many of their predicted receptors were specific to the ECM signaling network. Taken together, these pairs of transcription factors and receptors represent a number of signaling pathways which may be specifically activated in response to ECM (Fig. 5C). We show *Esrra* as a downstream target of *Il4ra* in myeloid cells in ECM, confirming alternative activation of macrophages. *Esrra* activates anti-inflammatory macrophages (Yuk et al., 2015) which are induced by IL4 and have been shown to be active in response to ECM. While there is nearly no detectable *Il4* in the single cell dataset (Supplementary Fig. 5B), we were able to detect signaling through *Il4ra* because we base signaling on activation of transcription factors. IL4 has been shown to be abundant in the ECM environment (Badylak and Gilbert, 2008) but is likely not detected in the single cell dataset due to dropout as the primary IL4 producers are eosinophils, T_H_2 cells, and potentially ILC2s (Heredia et al., 2013; Sadtler et al., 2016; Zhu, 2015).

We further show activation of *Sox17* in endothelial cells linked to expression of the receptor *Osmr* activated by *Osm* (Fig. 5C) from myeloid populations (Supplementary Fig. 5B). *Sox17* has been shown as critical for endothelial cell proliferation in response to injury (Liu et al., 2019) but has not been identified in ECM response. This signaling may be responsible for the increased proportion of endothelial cells in ECM and posits a mechanism for myeloid involvement in wound healing and a critical immune-tissue module communication. Finally, we also identify activation of *Ctcf*, a muscle-specific transcription factor (Delgado-Olguín et al., 2011), in fibroblasts linked to *Tnfrsf12a* or *TweakR*, recently shown to improve burn wound healing (Liu et al., 2018). The ligand for *TweakR, Tnfsf12* or *Tweak*, has elevated expression in both fibroblast and macrophage populations (Supplementary Fig. 5B). To our knowledge, *TweakR* has not been demonstrated as active in the ECM response. These three findings identify IL4, OSM, and TWEAK signaling pathways between fibroblasts and myeloid cells as potential therapeutic targets to promote regenerative wound healing and highlight the importance of immune-tissue module signaling.

The PCL specific transcription factors and their predicted receptors were also organized by enrichment in specific cell modules (Fig. 5E). In contrast to ECM, the PCL transcription factors associated with tissue-specific cell subsets had modest to low expression levels suggesting less activation of the tissue module (Fig. 5E). Instead, the tissue-specific module shared expression patterns similar to fibroblasts which may suggest suppression of tissue-specific regenerative programs. Using Domino’s signaling predictions, we identified a linkage between *Tgfbr2* and both *Sox11* and *Sox4* (Fig. 5F) in fibroblasts triggered by *Tgfb1* produced by myeloid populations (Supplementary Fig. 5C). Transforming growth factor beta (TGF-β) signaling is active in fibrotic environments like the foreign body response (Gerarduzzi and Di Battista, 2017). Bhattaram et al. demonstrated that *Sox4* and *Sox11* induce transformation of synoviocytes to fibroblast-like synoviocytes in rheumatoid arthritis (Bhattaram et al., 2018). These cells are thought to be responsible for the inflammatory environment that drives RA disease, but their existence and involvement in the biomaterials response was unknown. These findings suggest that induction of an inflammatory phenotype in fibroblasts driven by *Sox4* and *Sox11* may regulate TGF-β driven fibrosis. We also identify *Pirb* in the myeloid population correlated with pro-inflammatory transcription factor *Irf4*. While no readily accepted ligand for *Pirb* is known, it is an immune checkpoint regulating anti-inflammatory effects in myeloid populations (Bashirova et al., 2014). This suggests that *Pirb* may be a novel target to reduce chronic inflammation associated with fibrosis in response to biomaterials.

To test applicability of Domino in other data sets and diseases, we analyzed a publicly available dataset of healthy brain and brain with Alzheimer’s disease (AD) (Grubman et al., 2019). Both the AD and healthy signaling networks self-organized into modules associated with cell types (Supplementary Fig. 6A, D). The results for AD indicated *IL17RB* as a potential receptor upstream of *FOS*, which is active in AD (Anderson et al., 1994), and *NTRK3* upstream of *BHLHE40* in astrocytes (Supplementary Fig. 6B, C). *BHLHE40* has been linked with autoimmunity (Chih-Chung, 2017), connecting it with IL17 signaling (Zhu and Qian, 2012), which has been demonstrated as involved in AD (Cristiano et al., 2019). Domino also identified signaling pathways specific to healthy brain (Supplementary Fig. 6E). In astrocytes *NTRK2* targeted *PAX6*. Polymorphisms in *NTRK2* are correlated with AD (Zeng et al., 2013), and *PAX6* activates neurogenesis in astrocytes (Sakurai and Osumi, 2008) (Supplementary Fig. 6F). Finally, *SOX10*, a transcription factor promoting survival of myelin-producing oligodendrocytes (Takada et al., 2010), was activated by *ACVR1C* in healthy oligodendroctyes.

### Outlook

While the presented atlas represents a powerful resource for integration with future single cell data sets and investigation of biomaterials response, it is bound by the same limitations as single cell RNA sequencing. Only the top 10% of transcript from each cell is captured, making identification of more subtle changes in gene expression difficult and changes in lower expressed genes impossible. Cost limitations can also prevent higher numbers of biological replicates. Domino reconstructs signaling networks between modules of cells connected with transcriptional changes in cells independent of clustering analysis. Networks can be compared to identify signaling specific to conditions even with limited sample numbers or absence of key cell types. Domino was also able to capture cell interactions or influence with cells that were lost in scRNAseq (eosinophils) or where ligand expression was not captured with the limited depth of RNA sequencing in single cell. However, Domino is based solely on correlations and does not have rigorous statistical testing; therefore, it may be best suited for hypothesis generation with experimental validation.

Defining cell communication networks using Domino provides an unsupervised method to analyze single cell data sets that constructs and evaluates signaling networks using transcription factor activation. It generated signaling networks with modules connecting immune, stromal, and tissue-specific cell types independent of clustering information for both the biomaterial atlas. Comparison of networks yielded signaling pathways unique to experimental conditions, such as immune-tissue interactions in ECM, as well as tissue module suppression and inflammatory fibroblasts in the context of PCL. Communication between immune and tissue modules is an area of growing interest in tissue repair as demonstrated in immune-satellite signaling in muscle (Burzyn et al., 2013) and lymphatic-stem cell signaling in skin (Gur-Cohen et al., 2019). We believe Domino provides a new resource for unsupervised exploration of condition specific signaling patterns and generation of biologically testable signaling hypotheses that can be applied to an expanding biomaterials atlas and multiple other applications.

Overall, the biomaterial atlas describes new cell populations not previously defined in the biomaterial response, including NK cells and fibroblast subsets. The dataset and analysis methodology described here provides a cornerstone for future biomaterials development. Further expansion of the biomaterials atlas will enable comparison between a wide range of biomaterials implanted in different tissue environments and define cell signaling and communication patterns that regulate outcomes. This knowledge may facilitate better design of biomaterials to achieve desired responses such as tissue tolerance or repair and reduced incidence of adverse events associated with the foreign body response.

## Supporting information

All supplementary files

## Author Contributions

C.C. and J.H.E. conceptualized and drafted figures and manuscript, contributed to experimental design, and interpreted findings. C.C., J.H., J.A., and J.H.E. performed experiments and analyzed experimental results. C.C. wrote software. C.C., J.H.E., P.C., L.G., and E.F. contributed to computational methodology. All authors participated in construction of the manuscript and figures.

## Acknowledgements

The authors gratefully acknowledge financial support from the Morton Goldberg Chair, NIH Directors Pioneer Award, the Department of Defense, and Bloomberg-Kimmel Institute for Cancer Immunotherapy. Lana Garmire is supported by grants K01ES025434 awarded by NIEHS through funds provided by the trans-NIH Big Data to Knowledge (BD2K) initiative (www.bd2k.nih.gov), R01 LM012373 and LM012907 awarded by NLM, and R01 HD084633 awarded by NICHD. We thank Locke Davenport Huyer for help and expertise.

## Methods

### Volumetric muscle loss surgery

Animal procedures were performed in adherence to approved JHU IACUC protocols. Surgeries were performed when animals were 10 weeks of age. The bilateral traumatic muscle defect was created as previously described (Sicari et al., 2012). The defects were filled with 30 mg of a synthetic material or biological scaffold material. PCL was employed as a synthetic material (particulate, Mn = 50,000 g/mol, mean particle size < 600 μm, Polysciences). In turn, as a biological scaffold material, decellularized urinary bladder matrix (Matristem, Acell) was implanted from 0.05 ml of a 400 mg/ml suspension in phosphate buffered saline (PBS). Control surgeries were injected with 0.05 ml of PBS as a no implant control. All materials were UV sterilized prior to use. Mice were given subcutaneous carprofen (Rimadyl, Zoetis) at 5 mg/kg for pain relief. For the sample harvest, mice were euthanized at 1 and 6 weeks post-surgery.

### Flow Cytometry

Tissue samples were obtained by cutting the quadriceps femoris muscle from the hip to just above the knee. Tissues were finely diced manually with standard razor blades and digested for 45 min at 37°C with 1.67 Wünsch U/mL (0.5 mg/mL) Liberase TL (Roche Diagnostics, Sigma Aldrich) and 0.2 mg/mL DNase I (Roche Diagnostics, Sigma Aldrich) in RPMI-1640 medium supplemented with L-Glutamine and 15 mM HEPES (Gibco). The digested tissues were then pestled through 70 μm cell strainers (ThermoFisher Scientific) with RPMI-1640 (supplemented as before), and then washed twice with 1X DPBS. A Percoll (GE Healthcare) density gradient centrifugation was used to enrich the leukocyte fraction and remove blood and debris from the muscle samples and centrifuged at 2100 x g for 30 min with the lowest acceleration and no brake. Cells were washed 1X DPBS, stained with a viability live/dead amine reactive dye, washed with 1X DPS, blocked with anti-mouse TruStain FcX (BioLegend), and surface stained. Flow cytometry was performed using an Attune NxT Flow Cytometer (ThermoFisher Scientific) with a Violet, Blue, Green, and Red laser configuration. All subsequent analyses were performed on Live singlet cells using Windows based FlowJo^™^ software v10 (Benton Dickinson), license supplemented courtesy of the Johns Hopkins Bloomberg Flow Cytometry and Immunology Core.

| Mouse mAb | Fluorophore | Clone | Dilution | Stain Time | Volume |
| --- | --- | --- | --- | --- | --- |
| CD19 | PerCp-Cy5.5 | 6D5 | 1:300 | 45 mins | 50 µL |
| CD4 | PE | GK1.5 | 1:400 | 45 mins | 50 µL |
| SiglecF | PE-CF594 | E50-2440 | 1:400 | 45 mins | 50 µL |
| F4/80 | PE-Cy7 | EMR1 | 1:400 | 45 mins | 50 µL |
| Ly6g | APC | 1A8 | 1:250 | 45 mins | 50 µL |
| CD3 | AF700 | 500A2 | 1:250 | 45 mins | 50 µL |
| CD8a | APC-Cy7 | 53-6.7 | 1:250 | 45 mins | 50 µL |
| Ly6c | BV421 | HK1.4 | 1:250 | 45 mins | 50 µL |
| CD45 | BV510 | 30-F11 | 1:150 | 45 mins | 50 µL |
| Live Dead | Fixable Yellow | - | 1:1000 | 30 mins | 100 µL |
| CD11b | BV711 | M1/70 | 1:250 | 45 mins | 50 µL |

### NanoString Gene Expression Analysis

Muscles harvested from animals treated with ECM or PCL were immediately submerged in FisherSci TRIzol™ Reagent followed tissue mincing and grinding. Qiagen RNeasy Mini Kits were used to purify RNA following TRIzol RNA extraction. Purified RNA was quantified using an Agilent 2100 Bioanalyzer RNA 6000 Nano Kit according to manufacturer’s protocol. All RNA samples had RNA integrity score (RIN score) greater than 8.0. The NanoString™ nCounter© system was used with the nCounter© Fibrosis Gene Expression Panel following manufacturer’s recommendations with three biological replicates of each condition. Finally, the nSolver© software suite was used to analyze gene expression counts using the Advanced Analysis 2.0 module with default QC settings comparing the ECM and PCL samples with saline samples as a reference. To determine genes specific to ECM or PCL, we removed statistically significant genes in both ECM and PCL with the same direction of fold change. Volcano plots were generated using the EnhancedVolcano R package.

### Histological staining

After harvest, muscle samples were fixed in 10% neutral buffered formalin for 48 hours followed by ethanol and xylenes dehydration and paraffin embedding. Sections were rehydrated by ethanol gradient before staining. For hematoxylin and eosin staining, slides were stained in hematoxylin for 10m and eosin for 3m. We used the Masson’s Trichrome Stain Kit (Sigma Aldrich) following manufacturer’s guidelines. In both cases, slides were dehydrated by ethanol gradient, mounted using Fisher Chemical Permount Mounting Medium, and imaged with a Zeiss Axio Observer with Apotome.2 (Zeiss) with Zeiss Zen Blue software ver. 2.5 (Zeiss).

### Data and code availability

All raw data, processed files, and commented code are made available at GEOXXXXXXXXXX. Domino, our software package for analysis of intercellular communication, and sctools, a set of functions used in our code, are available at github.com/chris-cherry.

### Data preprocessing

Seurat was used for most processing steps where other software is not specified (Satija et al., 2015). All cell counts were pruned of cells with UMI counts below 250, cells with more than 10% mitochondrial genes, and genes expressed in fewer than 0.1% of cells. We then normalized and scaled the data with regression on UMI count and percent mitochondrial genes and calculated principle components using the top 2000 most variable genes. For muscle data sets we then corrected the principle components for batch effect using Harmony (Korsunsky et al., 2019). UMAP and shared nearest neighbor graph construction with subsequent Louvain clustering was then run on principle components.

### Cluster composition by condition

To calculate contribution of condition by cluster, we used only the CD45^+^ enriched samples to avoid biases in clusters containing sorted cells (i.e. fibroblasts and macrophages). Clusters from CD45^+^ and CD45^-^ cells were normalized separately to avoid slight differences in percent of CD45^+^ cells from enrichment by sample skewing normalization. For each sample, total number of cells by cluster were calculated and then normalized to the total of CD45^+^ or CD45^-^ cells in the dataset for the sample. The proportions of each sample were then averaged by condition to determine a condition-level average. Finally, in order to calculate a projected cell number, calculated proportions by sample (% of CD45^+^ cells) were multiplied by the total number of CD45^+^ cells determined by flow cytometry. CD45^-^ cells were not calculated due to inclusion of large amounts of debris in flow cytometry data without a positive marker.

### Phenotypic assignment of clusters

Differential expression testing for clusters was run using Mann-Whitney U tests. Each cluster was compared against all other clusters. The resulting gene expression profiles were examined to determine cluster phenotype. In many cases, unique expression of marker genes was sufficient to determine cluster identity. For the fibroblast clusters, we additionally used CytoSCAPE to determine stemness and assign precursor clusters. Finally, we cross-referenced the source publication for the macrophage dataset to determine precursor macrophage clusters (Sommerfeld et al., 2019).

### Gene regulatory network analysis

We used the SCENIC (Aibar et al., 2017) analysis pipeline to identify modules of genes targeted by transcription factors and calculate cell level enrichment scores. Genome ranking databases and *cis-*regulatory motif annotations were obtained from cisTarget Databases. First, we used Arboreto to fit a stochastic gradient boost machine using transcription factor counts to predict gene counts. Modules of genes targeted by transcription factor were then formed from the adjacencies, including genes with feature importances greater than the 95 percentile. The modules were then pruned, cross-referencing the motif annotations and ranking databases to remove modules with less than 80% of genes mapping to regions near binding sites for transcription factors or with less than 20 gene targets. Finally, enrichment for these modules was calculated using AUCell to identify cells with enrichment of genes targeted by transcription factors.

### Construction of a global signaling network

A list of human ligands, receptors, and their signaling relationships was obtained from CellphoneDB2 (Efremova et al., 2020). We then used biomaRt (Durinck et al., 2009) to convert genes from HGNC to MGI symbols, taking all conversions for each gene when multiple were found. Prior to calculating signaling relationships, counts matrices were pruned for genes expressed in fewer than 2.5% of cells. Pearson correlations were calculated between transcription factor activation scores and normalized, z-scored expression for identified receptors across all cells. Correlation between receptors which were determined as transcription factor targets by gene regulatory network analysis were then set to zero. This prevents targets of transcription factors which would be correlated with transcription factor activation from being interpreted as upstream of its transcription factor. Finally, transcription factors and receptors were considered signaling connections if Pearson correlation was greater than 0.3 with a maximum of ten receptors per transcription factor. For all receptors connected with transcription factors, ligand signaling partners were identified from the CellphoneDB2 database. Ligands that were not found in the dataset were excluded.

### Comparison of condition specific global signaling networks

In order to identify intracellular signaling patterns associated with an independent variable, we first split the single cell data by that variable. More specifically, we split the volumetric muscle dataset by treatment with ECM or PCL and the Alzheimer’s dataset by disease status of the patients. We then constructed a global signaling network for each of the separate data sets with identical parameters. Finally, we identified network items specific to each condition by set subtraction. Transcription factors or receptors only present in one condition’s signaling network were considered as potentially condition specific.

### Cluster specific subnetwork identification

In order to identify intracellular signaling patterns within a cluster with ligands responsible for their activation, we first identified active transcription factors by cluster using Mann-Whitney U tests. For each cluster, the top over-expressed genes were selected based on p-value with positive log fold change as compared to all other clusters. Transcription factors with p-values below .001 were included with up to 10 transcription factors per cluster. We then generated a signaling subnetwork for each cluster by pruning all network items not connected to the cluster enriched transcription factors.

### Prediction of intercellular signaling networks

Using the cluster specific signaling subnetworks, we identified ligands most likely to be responsible for activation of cellular phenotype for each target cluster. It’s important to note that in the data sets we’ve analyzed there tend to be many ligands below detection threshold, so we don’t use signaling pathways without expressed ligands in construction of intercellular signaling networks. To calculate intercluster signaling with ligands found in the data set, we first averaged z-scored ligand expression by cluster. We then generated a signaling score by summing the averaged z-scores by cluster. The signaling represents whether a particular cluster is over- or under-expressing the ligands predicted to activate the target cluster. These values are used as directed, weighted edges between clusters as nodes to construct an intercellular signaling network.

---

**Supplementary Figure 1:**
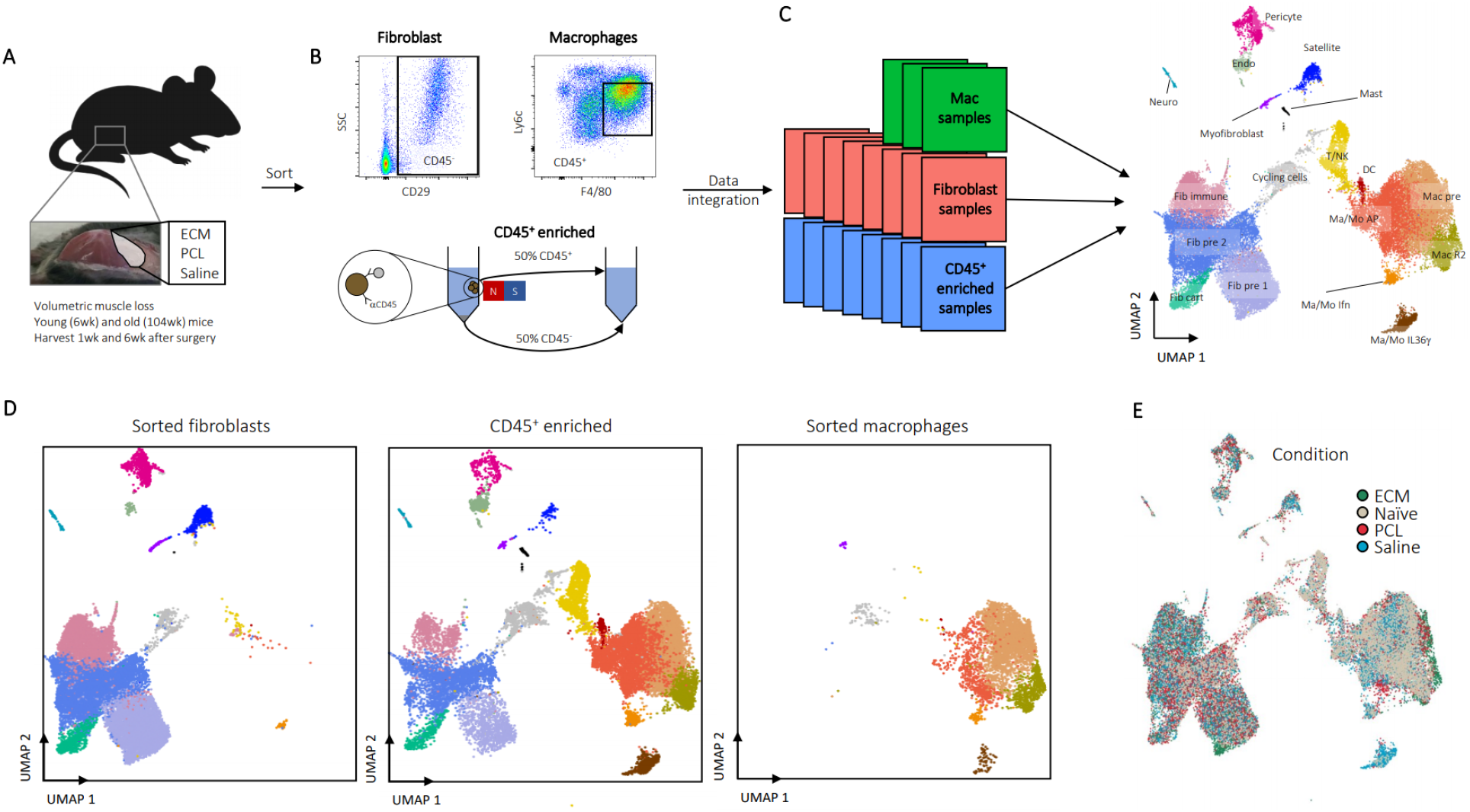
Experimental Overview of Assembled Data Sets. **A**, All data sets were taken from mice after volumetric muscle loss treatment. After surgical excision of a large portion of the quadriceps, the wound site was filled with a biomaterial or saline control and stapled shut. Mice were then harvested 1 or 6 weeks after surgery. Young (6 week) or aged (104 week) old animals were used. **B**, At time of harvest, cells were isolated one of three ways after digestions. For macrophages, cells were sorted as CD45^+^F4/80^Hi^Ly6c^+^, for fibroblasts cells were sorted as CD45^-^CD19^-^CD29^+^, and for the all-cell dataset CD45^+^ cells were enriched to ~50% using MACS beads. **C**, Data sets were integrated for analysis using Harmony. A complete summary of available data sets is given in Supplementary Table 2. **D**, Enrichment of fibroblasts and macrophages due to inclusion of sorted fibroblast and macrophage data sets. The sorted fibroblasts (left) and macrophages (right) are shown in comparison to the CD45^+^ enriched sample (middle). **E**, Cells by condition. Cells are colored by condition plotted on UMAP dimensions. Cells were plotted in order of ECM, PCL, Saline, and Naïve.

**Supplementary Figure 2:**
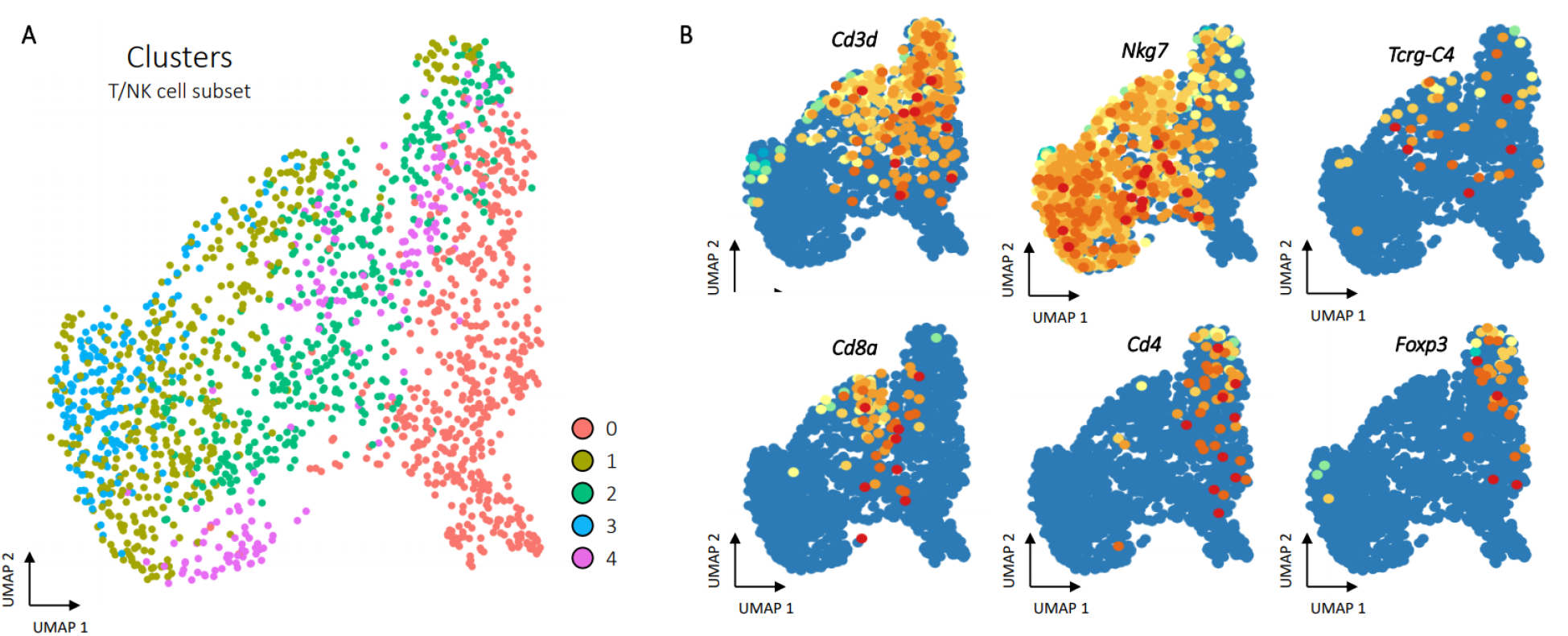
Subset clustering of T/NK cells. **A**, Clustering of T/NK cells. After subsetting to only T/NK cells, principle component analysis, clustering, and UMAP was run following the same procedures as the whole dataset. The five resulting clusters are visualized in the T/NK cell specific UMAP space. **B**, Gene expression for T and NK cell markers. Gene expression for T cell markers *Cd3d* (all T cells), *Cd8a* (Th1 cells), *Cd4* (Th2 cells), and *Tcrg-C4* (γδ T cells), as well as NK cell marker *Nkg7* are shown.

**Supplementary Figure 3:**
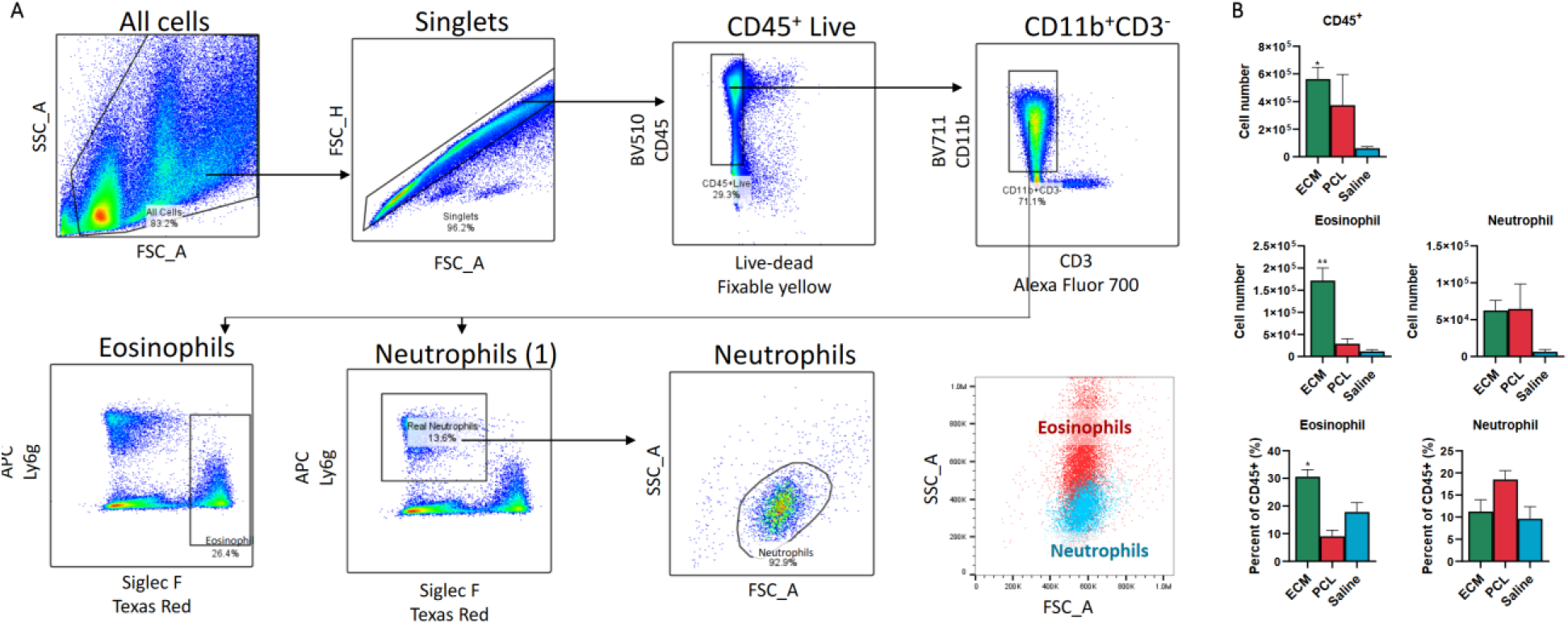
Flow cytometry of neutrophils and eosinophils. **A**, Cells were gated on scatter (FSC_A, SSC_A) followed by doublet discrimination (FSC_A, FSC_H), selection of CD45^+^ live cells (Fixable Yellow-, CD45+), and selection of myeloid cells (CD3-, CD11b+). This population was used to identify eosinophils (Ly6g low, Siglec F+) and neutrophils (Ly6g+, Siglec F-) of correct size (FSC_A, SSC_A). **B**, Numbers of CD45^+^ cells, eosinophils, and neutrophils from ECM, PCL, or saline treated animals one week after surgery (top). Eosinophil and neutrophil amounts as proportion of CD45^+^ cells are given below. Data are mean ± SEM. \*\**P* < 0.01, \**P* < 0.05 by analysis of variance (ANOVA) followed by Dunnett’s multiple comparison testing where *P* is adjusted p value.

**Supplementary Figure 4:**
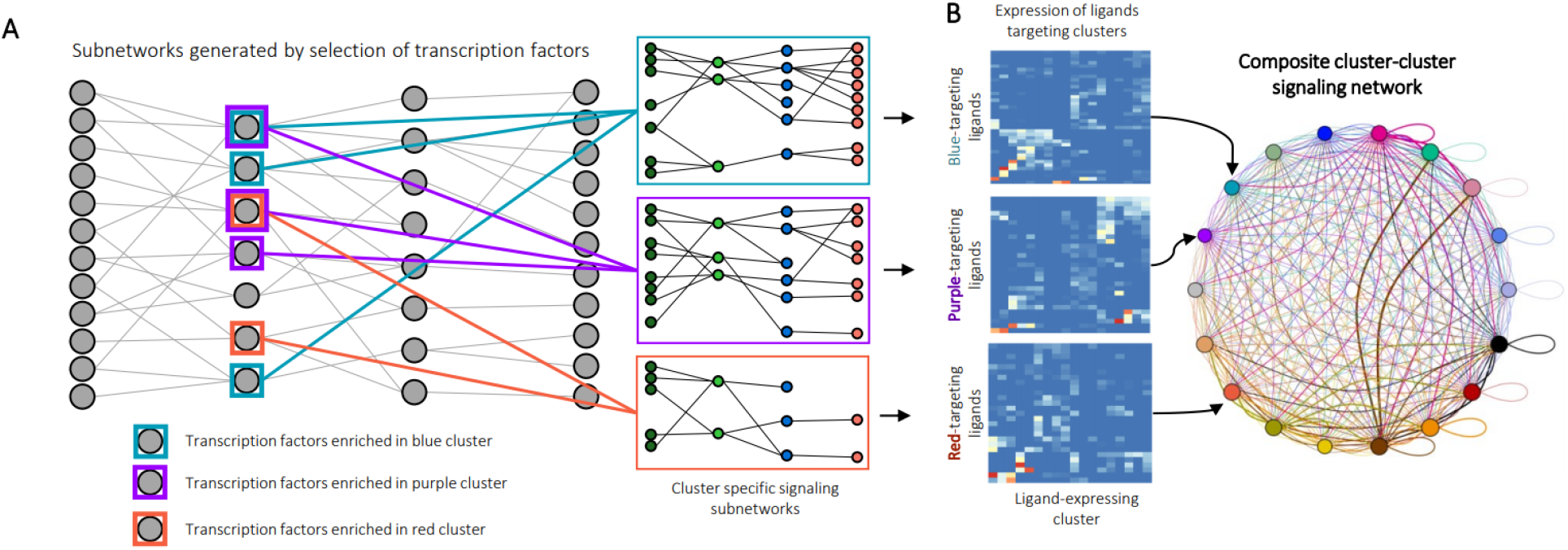
Identification of intercluster signaling with Domino. **A**, Identification of cluster-specific signaling subnetworks. Transcription factors enriched by cluster are identified by Wilcoxon rank sum and networks pruned for disconnected nodes to generate signaling subnetworks relevant for biological activation of clusters. **B**, Calculation of intercluster signaling networks. Once phenotypically relevant receptors are identified by cluster specific signaling subnetworks, cluster-cluster signaling scores are calculated by cluster averaged scaled expression of ligands present in cluster-specific subnetworks. Every potential cluster-cluster combination is scored, and these weights used to generate an intercluster signaling network.

**Supplementary Figure 5:**
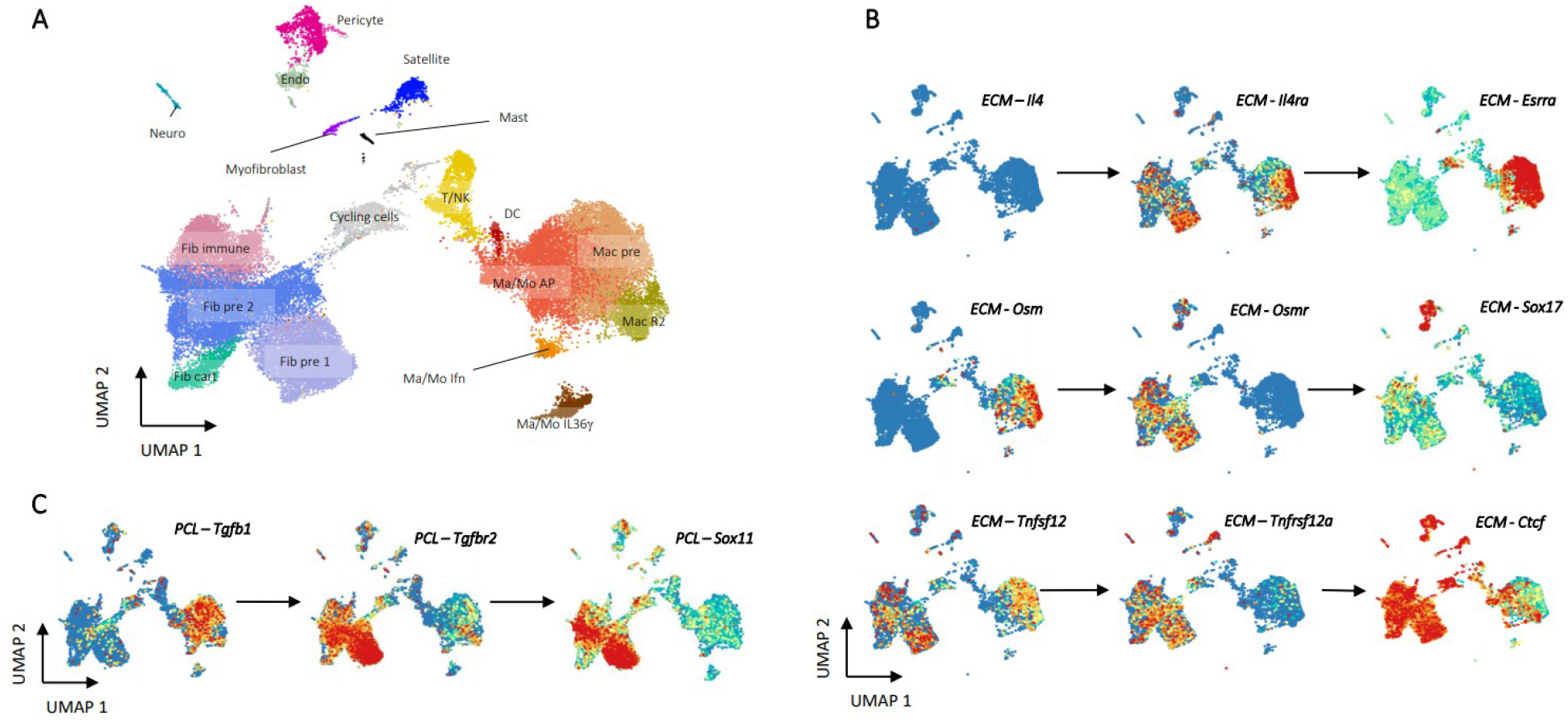
Signaling pathways for ECM and PCL treated cells visualized with their known ligands. **A**, A UMAP plot with clusters labeled for reference when viewing feature plots. **B**, ECM specific signaling pathways specified in Figure 3. Each pathway contains gene expression of ligands and receptors from each pathway, as well as transcription factor activation scores for the predicted transcription factor target. Ligands completely absent from the data set are not shown, although they may still be viable targets for a target receptor. **C**, PCL specific signaling pathways specified in Figure 3. No readily accepted ligands for *Pirb* have been identified, so the *Pirb Irf4* pathway is not shown.

**Supplementary Figure 6:**
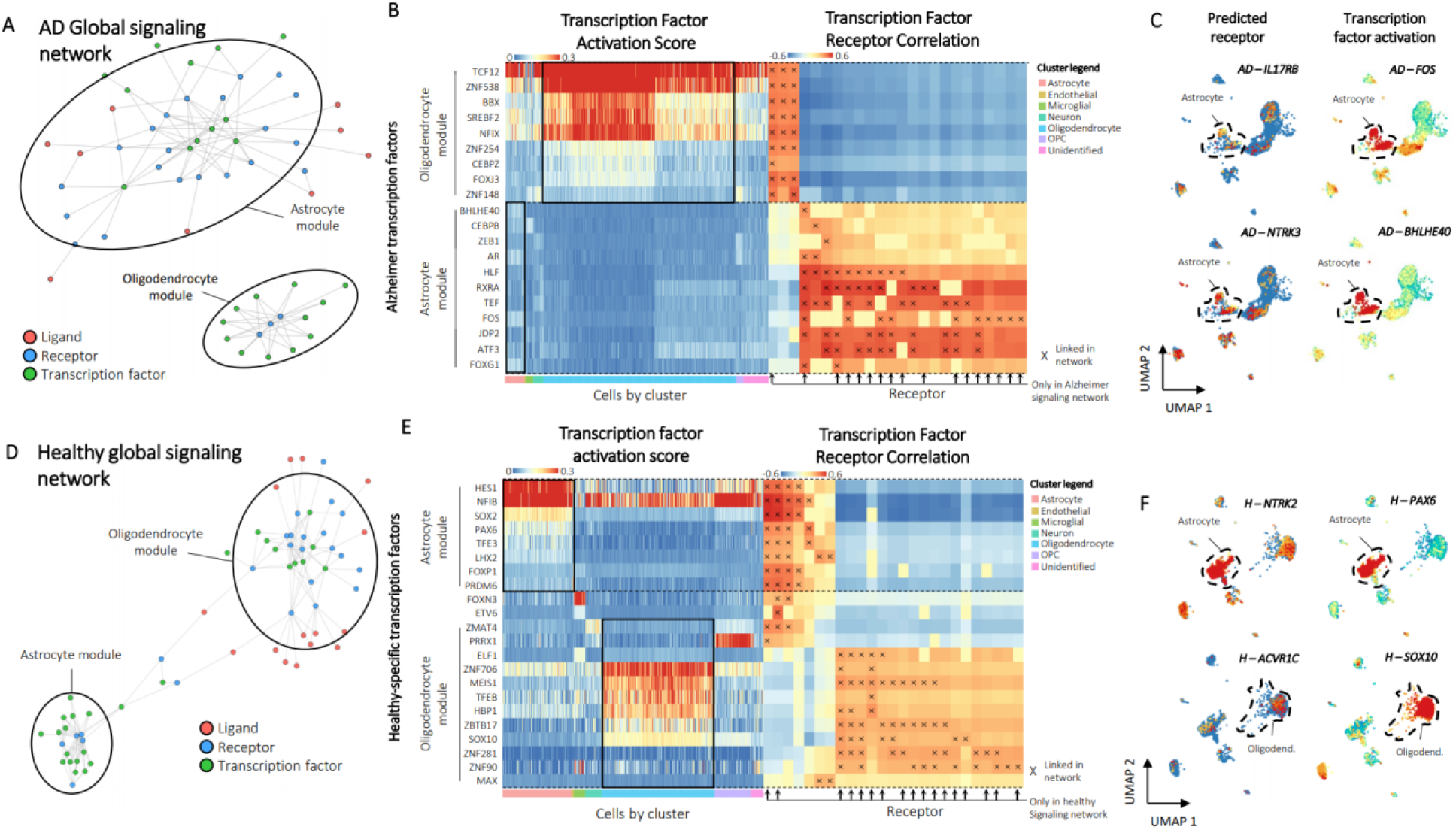
Pathological signaling found from a public dataset of Alzheimer’s Disease. **A**, The Alzheimer’s disease (AD) global signaling network. two modules of receptors and transcription factors are readily apparent and labeled based on enrichment of transcription factors by cluster. **B**, Heatmaps of transcription factor activation score for AD-specific transcription factors (left) and correlation of transcription factor activation score with receptor expression (right). Transcription factors are binned according to their membership to the astrocyte or oligodendrocyte modules from the AD global signaling network. Cells are ordered and colored according to their cluster. Receptors found only in AD are marked with arrows. Connections between receptor and transcription factors are marked with an ‘x’ on the correlation heatmap. **C**, Example feature plots of gene expression and activation scores for specific receptor-transcription factor pairs identified by domino in the AD condition. **D**, The healthy global signaling network. Two modules of receptors and transcription factors are readily apparent and labeled based on enrichment of transcription factors by cluster. **E**, Heatmaps of transcription factor activation score for healthy-specific transcription factors (left) and correlation of transcription factor activation score with receptor expression (right). Transcription factors are binned according to their membership to the astrocyte or oligodendrocyte modules from the global signaling network. Cells are ordered and colored according to their cluster. Receptors found only in the healthy signaling network are marked with arrows. Connections between receptor and transcription factors are marked with an ‘x’ on the correlation heatmap. **F**, Example feature plots of gene expression and activation scores for specific receptor-transcription factor pairs identified by domino in healthy cells.

## Notes

### Competing Interest Statement

The authors have declared no competing interest.

