## Supplementary material for "Intercellular signaling dynamics from a single cell atlas of the biomaterials response": All supplementary files: Hires Figures.pdf

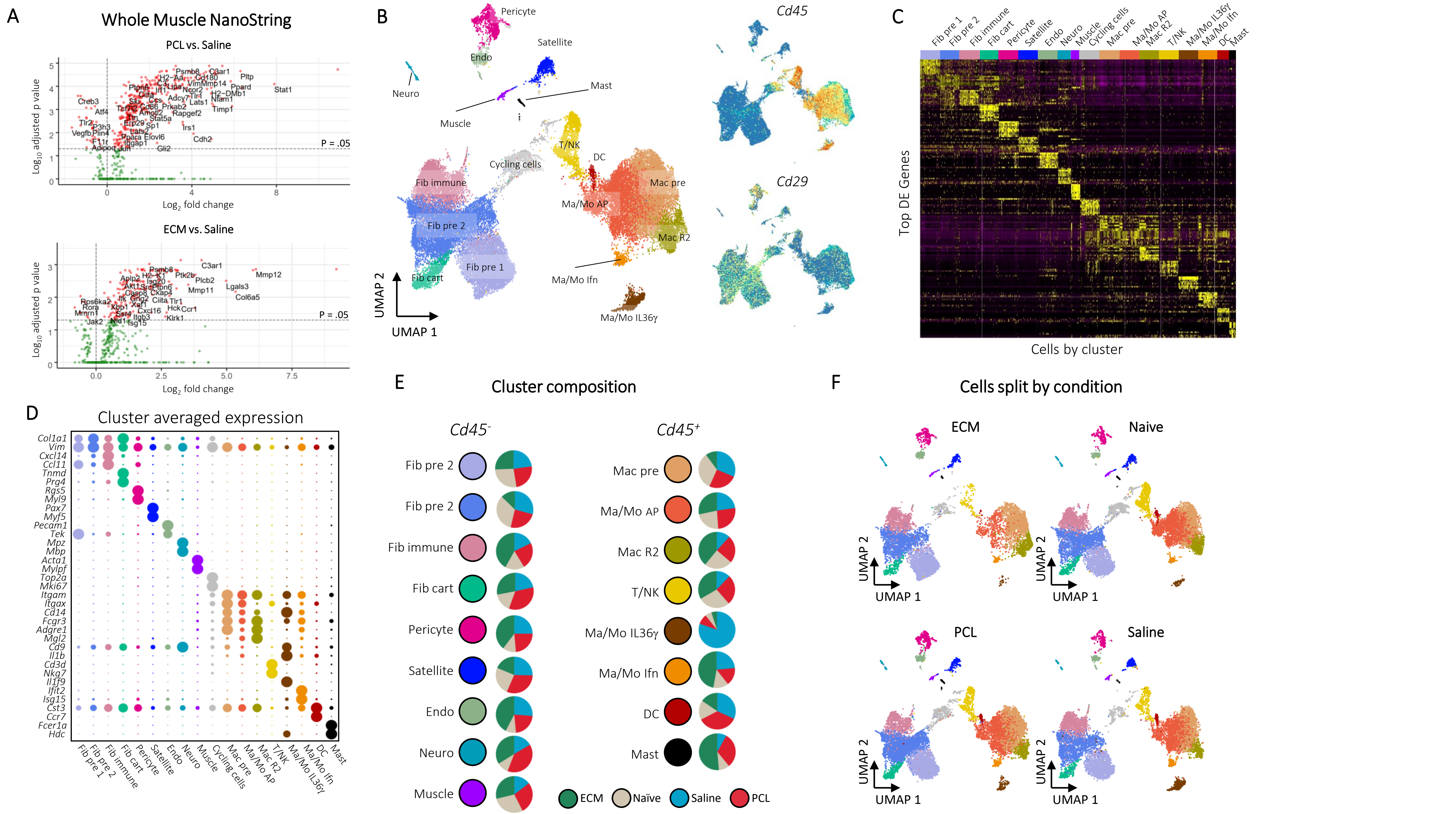
A
Whole Muscle NanoString
PCL vs. Saline
ECM vs. Saline
Log10 adjusted p value
Log2 fold change
P = .05
B
Cd45
Cd29
UMAP 1
UMAP 2
C
Top DE Genes
Cells by cluster
D
Cluster averaged expression
E
Cluster composition
Cd45-
Cd45+
ECM
Naïve
Saline
PCL
F
Cells split by condition
ECM
Naive
PCL
Saline

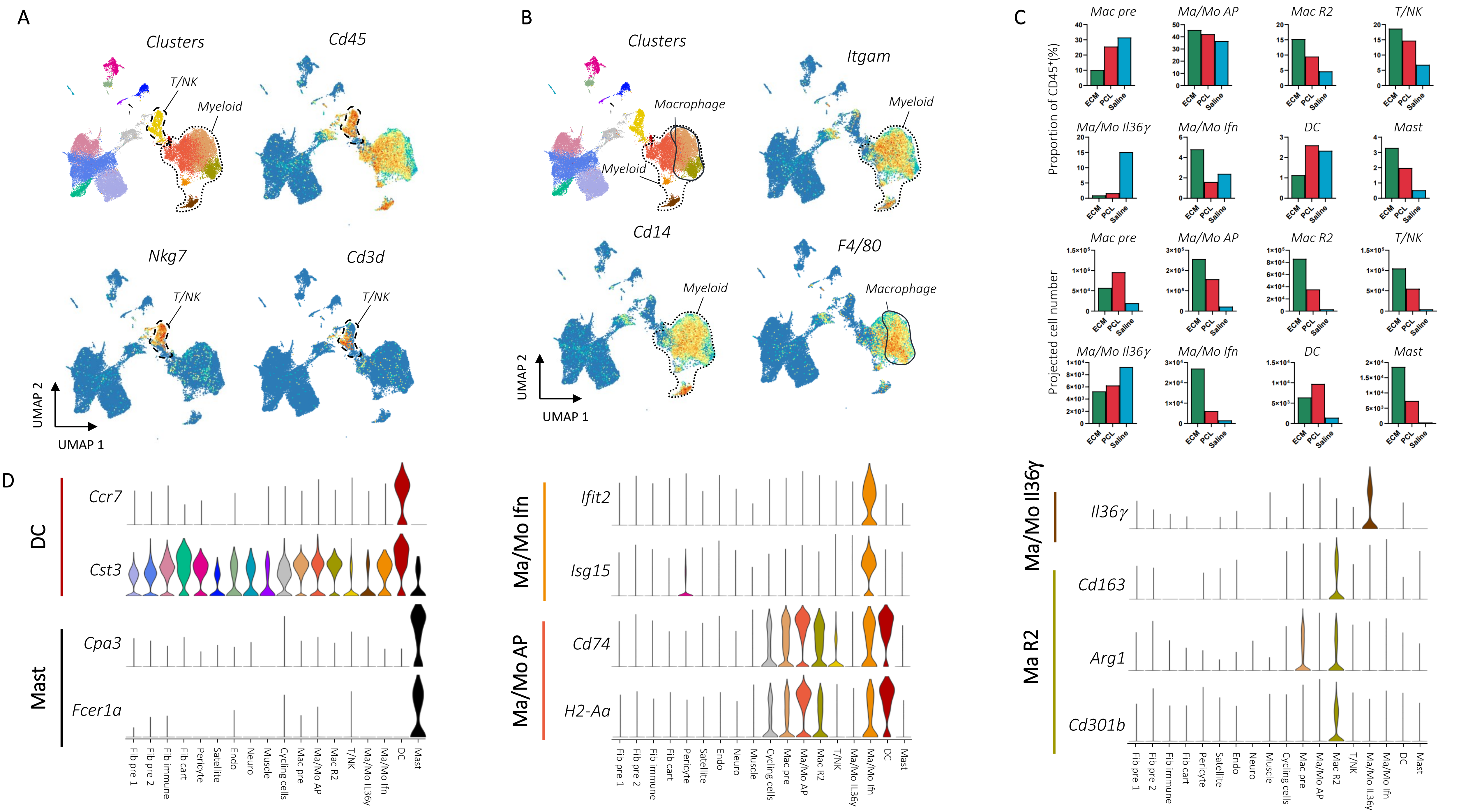

A
Clusters
Cd45
T/NK
Myeloid
Nkg7
Cd3d
UMAP 2
UMAP 1
B
Clusters
Macrophage
Myeloid
Itgam
Cd14
F4/80
C
Proportion of CD45+(%)
Projected cell number
Mac pre
Ma/Mo AP
Mac R2
T/NK
Ma/Mo Il36γ
Ma/Mo Ifn
DC
Mast
ECM
PCL
Saline
D
DC
Ccr7
Cst3
Mast
Cpa3
Fcer1a
Ma/Mo Ifn
Ifit2
Isg15
Ma/Mo AP
Cd74
H2-Aa
Ma/Mo Il36γ
Il36γ
Ma R2
Cd163
Arg1
Cd301b
Fib pre 1
Fib pre 2
Fib immune
Fib cart
Pericyte
Satellite
Endo
Neuro
Muscle
Cycling cells
Mac pre
Ma/Mo AP
Mac R2
T/NK
Ma/Mo Il36γ
Ma/Mo Ifn
DC
Mast

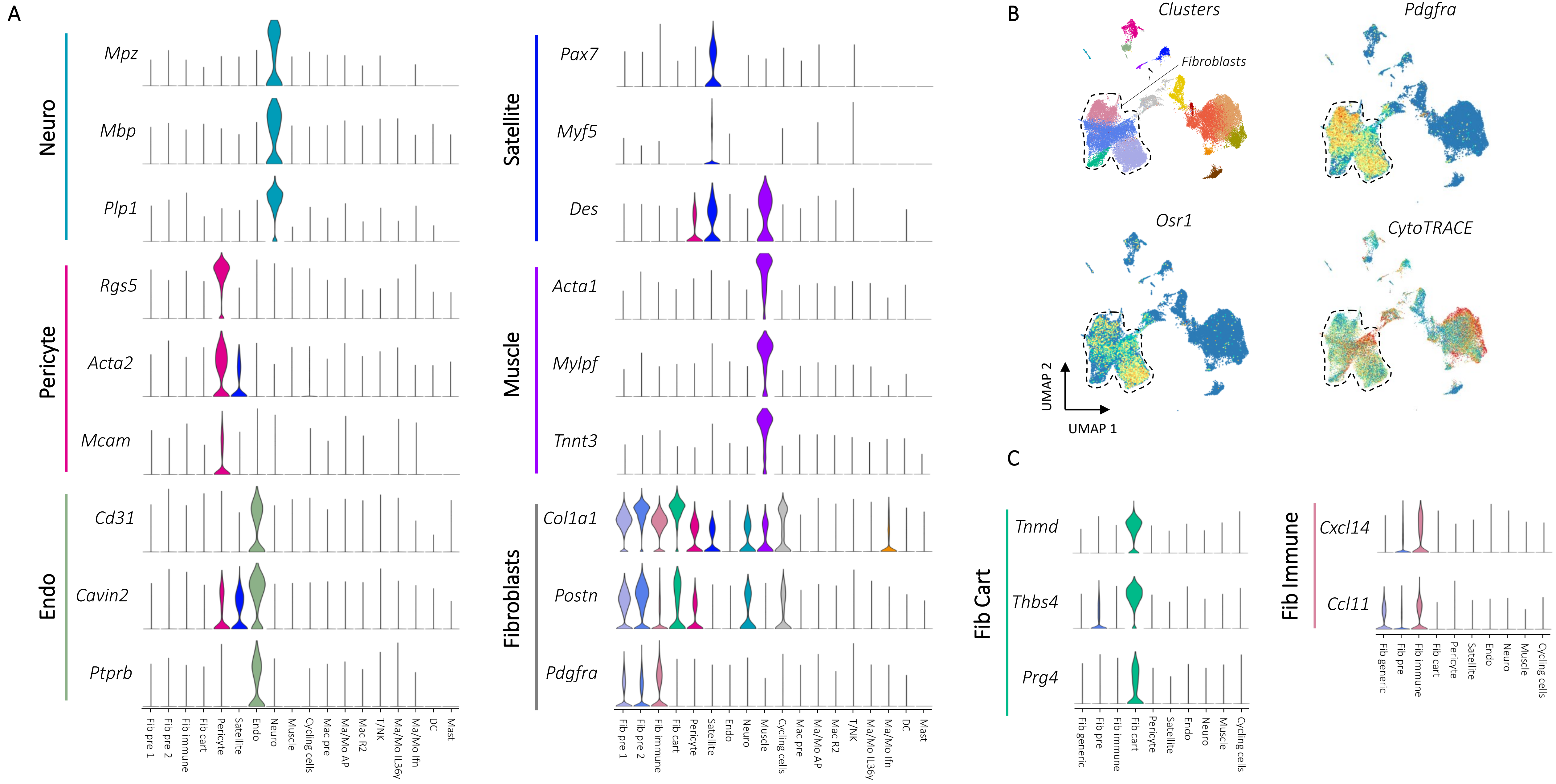

A
Neuro
Mpz
Mbp
Plp1
Pericyte
Rgs5
Acta2
Mcam
Endo
Cd31
Cavin2
Ptprb
Fib pre 1
Fib pre 2
Fib immune
Fib cart
Pericyte
Satellite
Endo
Neuro
Muscle
Cycling cells
Mac pre
Ma/Mo AP
Mac R2
T/NK
Ma/Mo Il36γ
Ma/Mo Ifn
DC
Mast
Satellite
Pax7
Myf5
Des
Muscle
Acta1
Mylpf
Tnnt3
Fibroblasts
Col1a1
Postn
Pdgfra
B
Clusters
Fibroblasts
Pdgfra
Osr1
CytoTRACE
UMAP 2
UMAP 1
C
Fib Cart
Tnmd
Thbs4
Prg4
Fib generic
Fib pre
Fib immune
Fib cart
Pericyte
Satellite
Endo
Neuro
Muscle
Cycling cells
Fib Immune
Cxcl14
Ccl11

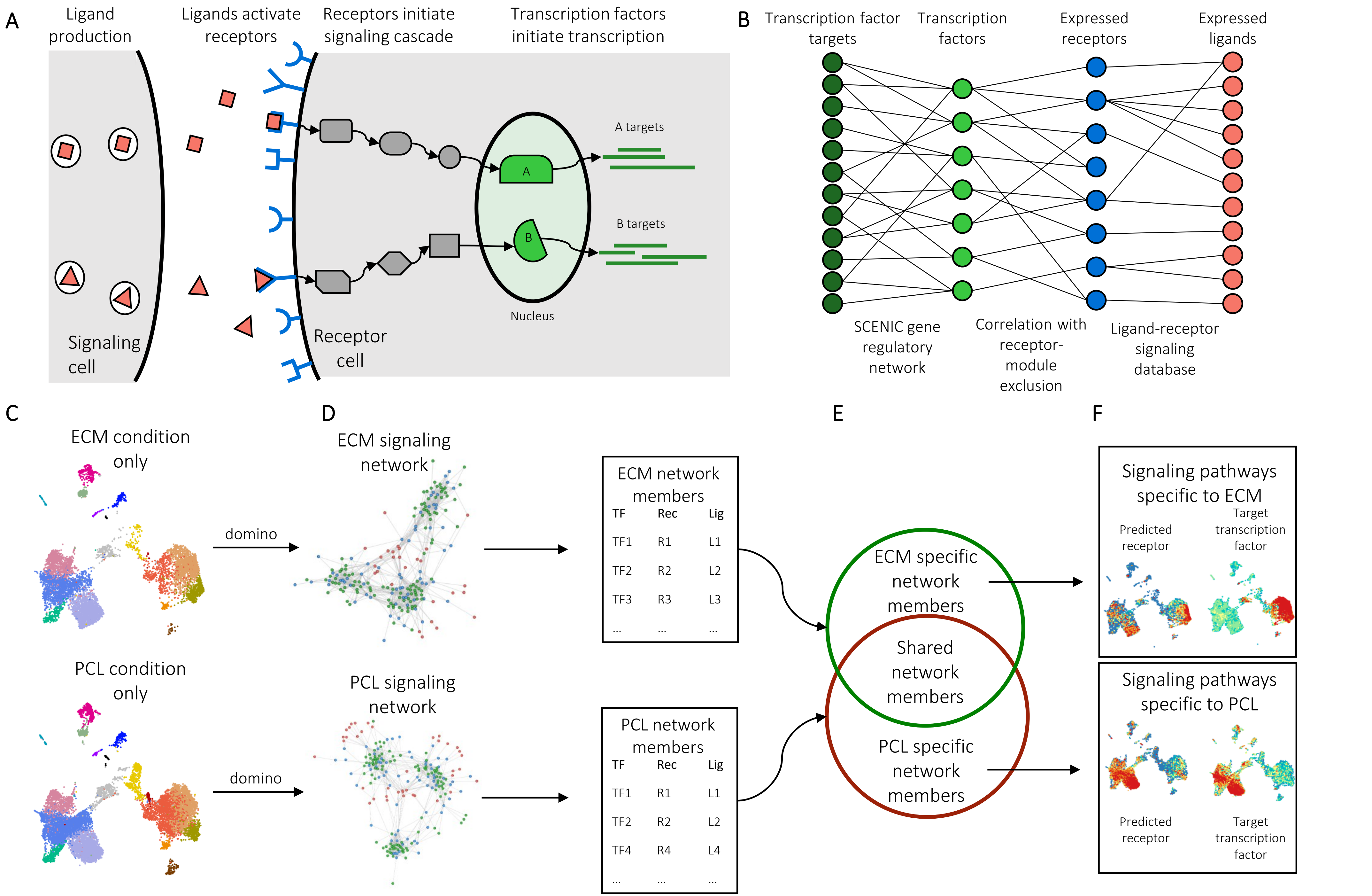
A
Ligand production
Ligands activate receptors
Receptors initiate signaling cascade
Transcription factors initiate transcription
A targets
A
B targets
B
Nucleus
Signaling cell
Receptor cell
B
Transcription factor targets
Transcription factors
Expressed receptors
Expressed ligands
SCENIC gene regulatory network
Correlation with receptor-module exclusion
Ligand-receptor signaling database
C
ECM condition only
PCL condition only
domino
domino
D
ECM signaling network
PCL signaling network
ECM network members
TF Rec Lig
TF1 R1 L1
TF2 R2 L2
TF3 R3 L3
... ... ...
PCL network members
TF Rec Lig
TF1 R1 L1
TF2 R2 L2
TF4 R4 L4
... ... ...
E
ECM specific network members
Shared network members
PCL specific network members
F
Signaling pathways specific to ECM
Predicted receptor
Target transcription factor
Signaling pathways specific to PCL
Predicted receptor
Target transcription factor

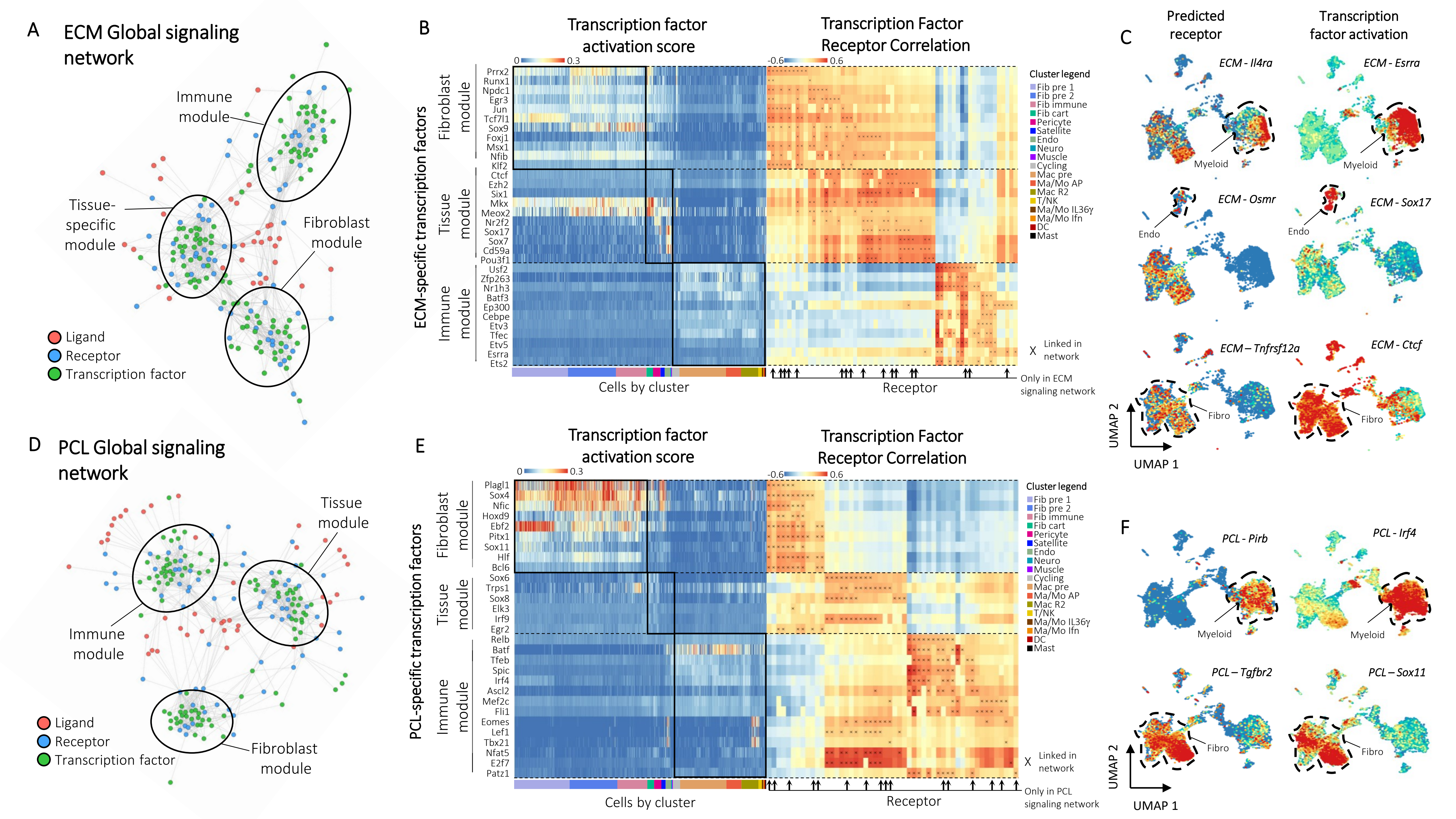
A
ECM Global signaling network
Immune module
Tissue-specific module
Fibroblast module
Ligand
Receptor
Transcription factor
B
Transcription factor activation score
Transcription Factor Receptor Correlation
ECM-specific transcription factors
Fibroblast module
Tissue module
Immune module
Prrx2
Runx1
Npdc1
Egr3
Jun
Tcf7l1
Sox9
Foxj1
Msx1
Nfib
Klf2
Ctcf
Ezh2
Six1
Mkx
Meox2
Nr2f2
Sox17
Sox7
Cd59a
Pou3f1
Usf2
Zfp263
Nr1h3
Batf3
Ep300
Cebpe
Etv3
Tfec
Etv5
Esrra
Ets2
Cells by cluster
Receptor
Cluster legend
Fib pre 1
Fib pre 2
Fib immune
Fib cart
Pericyte
Satellite
Endo
Neuro
Muscle
Cycling
Mac pre
Ma/Mo AP
Mac R2
T/NK
Ma/Mo IL36γ
Ma/Mo Ifn
DC
Mast
X Linked in network
Only in ECM signaling network
C
Predicted receptor
Transcription factor activation
ECM - Il4ra
ECM - Esrra
Myeloid
ECM - Osmr
ECM - Sox17
Endo
ECM – Tnfrsf12a
ECM - Ctcf
Fibro
UMAP 2
UMAP 1
D
PCL Global signaling network
Tissue module
Immune module
Fibroblast module
Ligand
Receptor
Transcription factor
E
Transcription factor activation score
Transcription Factor Receptor Correlation
PCL-specific transcription factors
Fibroblast module
Tissue module
Immune module
Plagl1
Sox4
Nfic
Hoxd9
Ebf2
Pitx1
Sox11
Hlf
Bcl6
Sox6
Trps1
Sox8
Elk3
Irf9
Egr2
Relb
Batf
Tfeb
Spic
Irf4
Ascl2
Mef2c
Fli1
Eomes
Lef1
Tbx21
Nfat5
E2f7
Patz1
Cells by cluster
Receptor
Cluster legend
Fib pre 1
Fib pre 2
Fib immune
Fib cart
Pericyte
Satellite
Endo
Neuro
Muscle
Cycling
Mac pre
Ma/Mo AP
Mac R2
T/NK
Ma/Mo IL36γ
Ma/Mo Ifn
DC
Mast
X Linked in network
Only in PCL signaling network
F
PCL - Pirb
PCL - Irf4
Myeloid
PCL – Tgfbr2
PCL – Sox11
Fibro
UMAP 2
UMAP 1

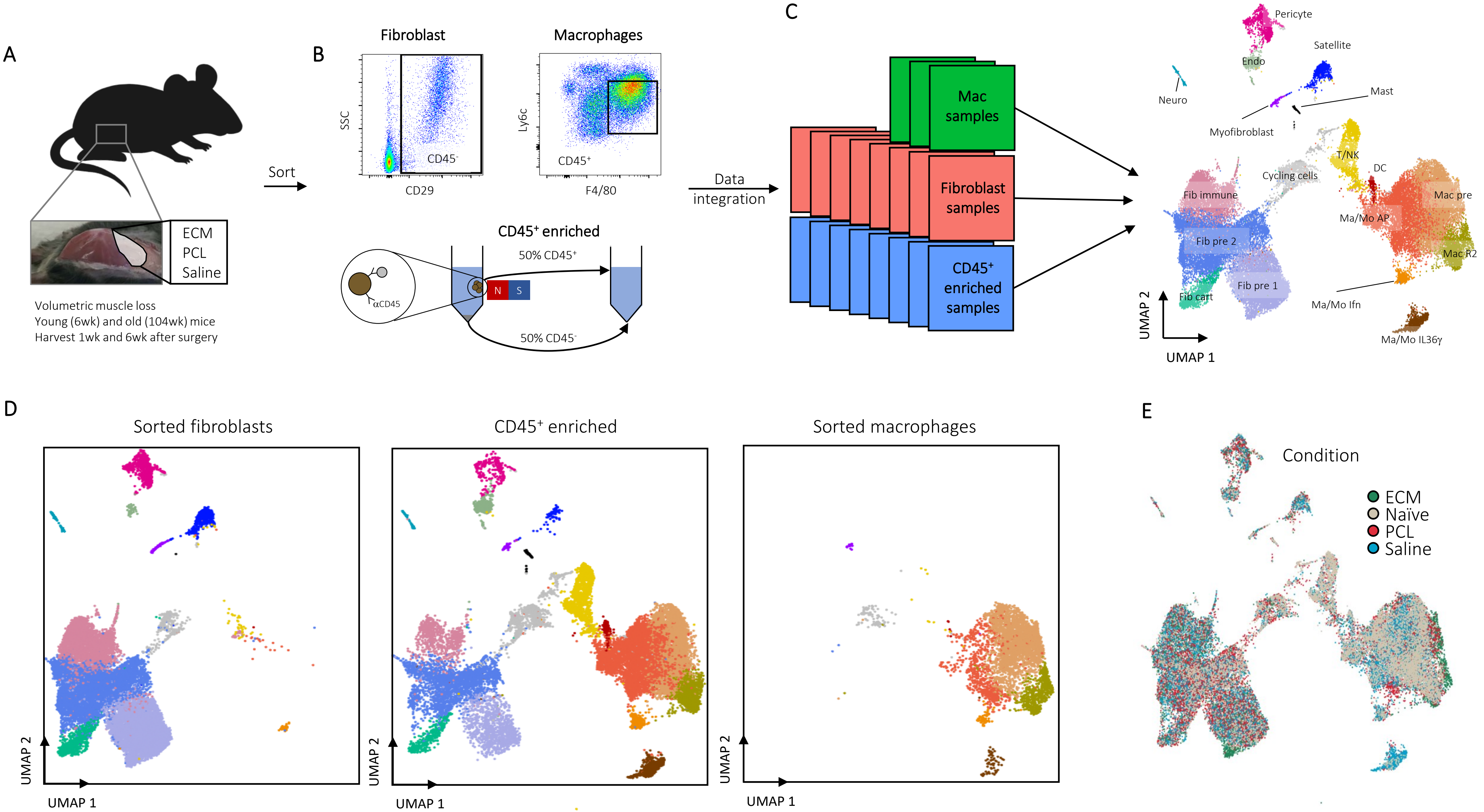
A
ECM
PCL
Saline
Volumetric muscle loss
Young (6wk) and old (104wk) mice
Harvest 1wk and 6wk after surgery
Sort
B
Fibroblast
Macrophages
SSC
CD29
CD45-
Ly6c
F4/80
CD45+
CD45+ enriched
αCD45
N
S
50% CD45+
50% CD45-
Data integration
C
Mac samples
Fibroblast samples
CD45+ enriched samples
Pericyte
Satellite
Endo
Neuro
Mast
Myofibroblast
T/NK
DC
Cycling cells
Fib immune
Mac pre
Ma/Mo AP
Fib pre 2
Mac R2
Fib pre 1
Fib cart
Ma/Mo Ifn
Ma/Mo IL36γ
UMAP 2
UMAP 1
D
Sorted fibroblasts
CD45+ enriched
Sorted macrophages
E
Condition
ECM
Naïve
PCL
Saline

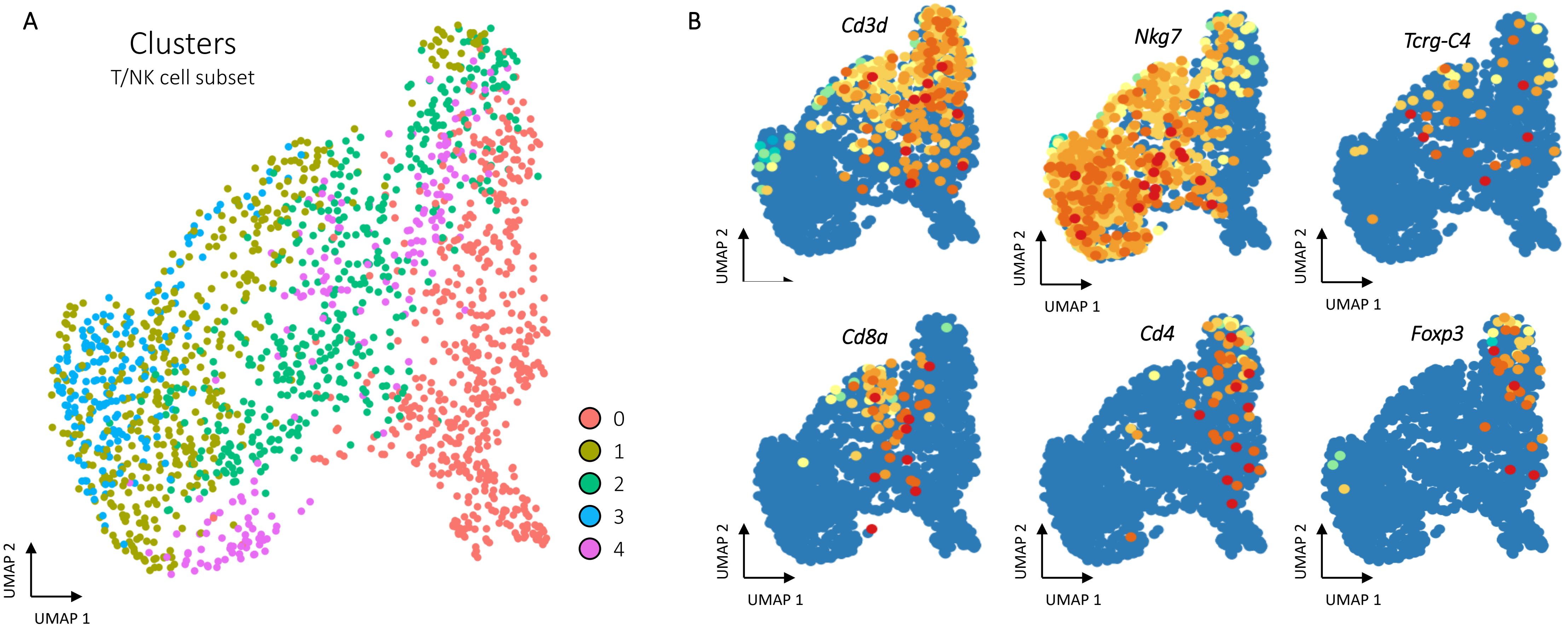

A
Clusters
T/NK cell subset
0
1
2
3
4
UMAP 2
UMAP 1
B
Cd3d
Nkg7
Tcrg-C4
Cd8a
Cd4
Foxp3
UMAP 2
UMAP 1

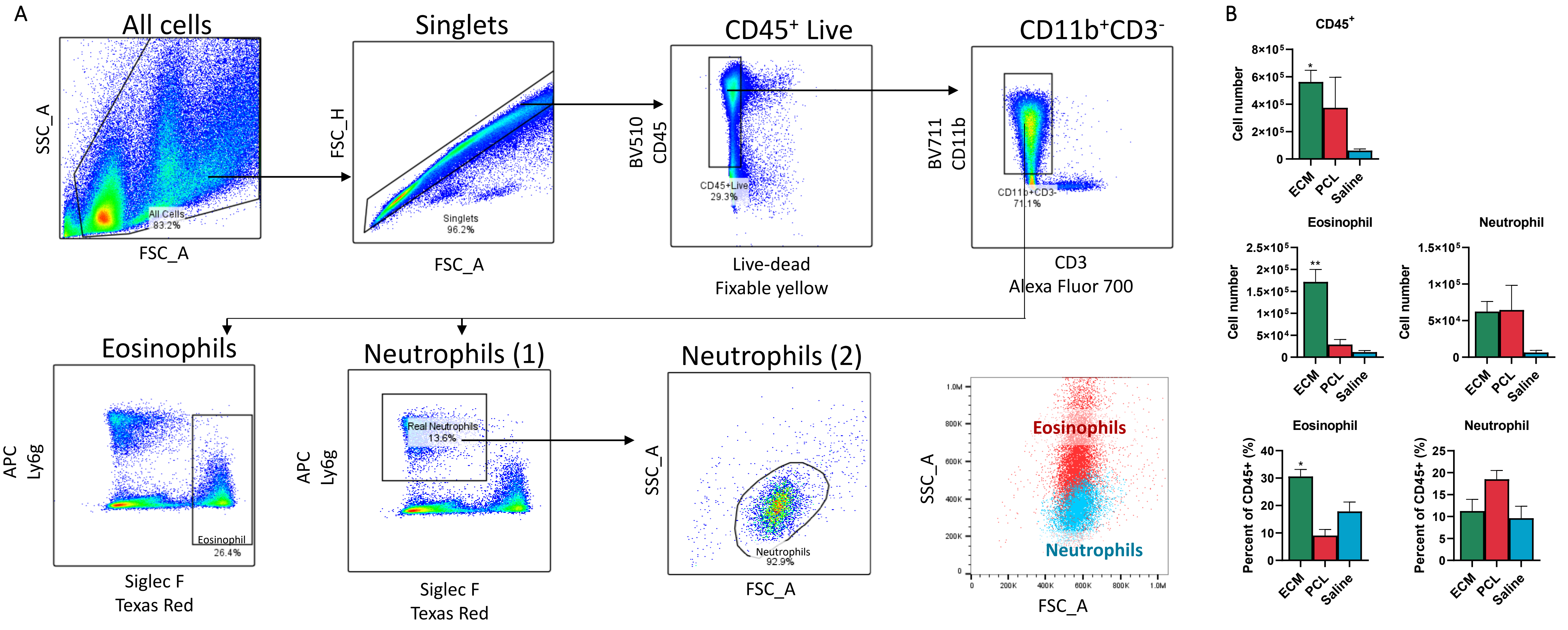

A
All cells
SSC_A
FSC_A
All Cells
83.2%
Singlets
FSC_H
Singlets
96.2%
FSC_A
CD45+ Live
BV510
CD45
CD45+Live
29.3%
Live-dead
Fixable yellow
CD11b+CD3-
BV711
CD11b
CD11b+CD3-
71.1%
CD3
Alexa Fluor 700
Eosinophils
APC
Ly6g
Eosinophil
26.4%
Siglec F
Texas Red
Neutrophils (1)
APC
Ly6g
Real Neutrophils
13.6%
Siglec F
Texas Red
Neutrophils (2)
SSC_A
Neutrophils
92.9%
FSC_A
SSC_A
Eosinophils
Neutrophils
FSC_A
B
CD45+
Cell number
ECM
PCL
Saline
Eosinophil
Neutrophil
Cell number
Eosinophil
Neutrophil
Percent of CD45+ (%)

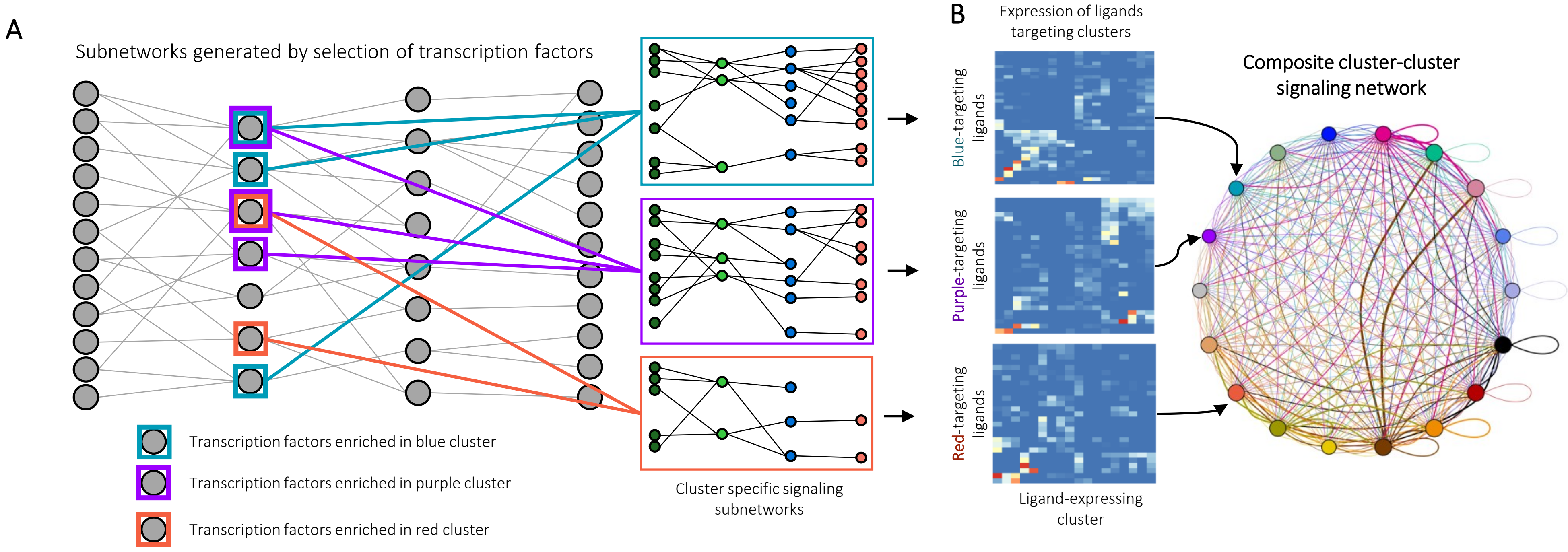
A
Subnetworks generated by selection of transcription factors
Transcription factors enriched in blue cluster
Transcription factors enriched in purple cluster
Transcription factors enriched in red cluster
Cluster specific signaling subnetworks
B
Expression of ligands targeting clusters
Blue-targeting ligands
Purple-targeting ligands
Red-targeting ligands
Ligand-expressing cluster
Composite cluster-cluster signaling network

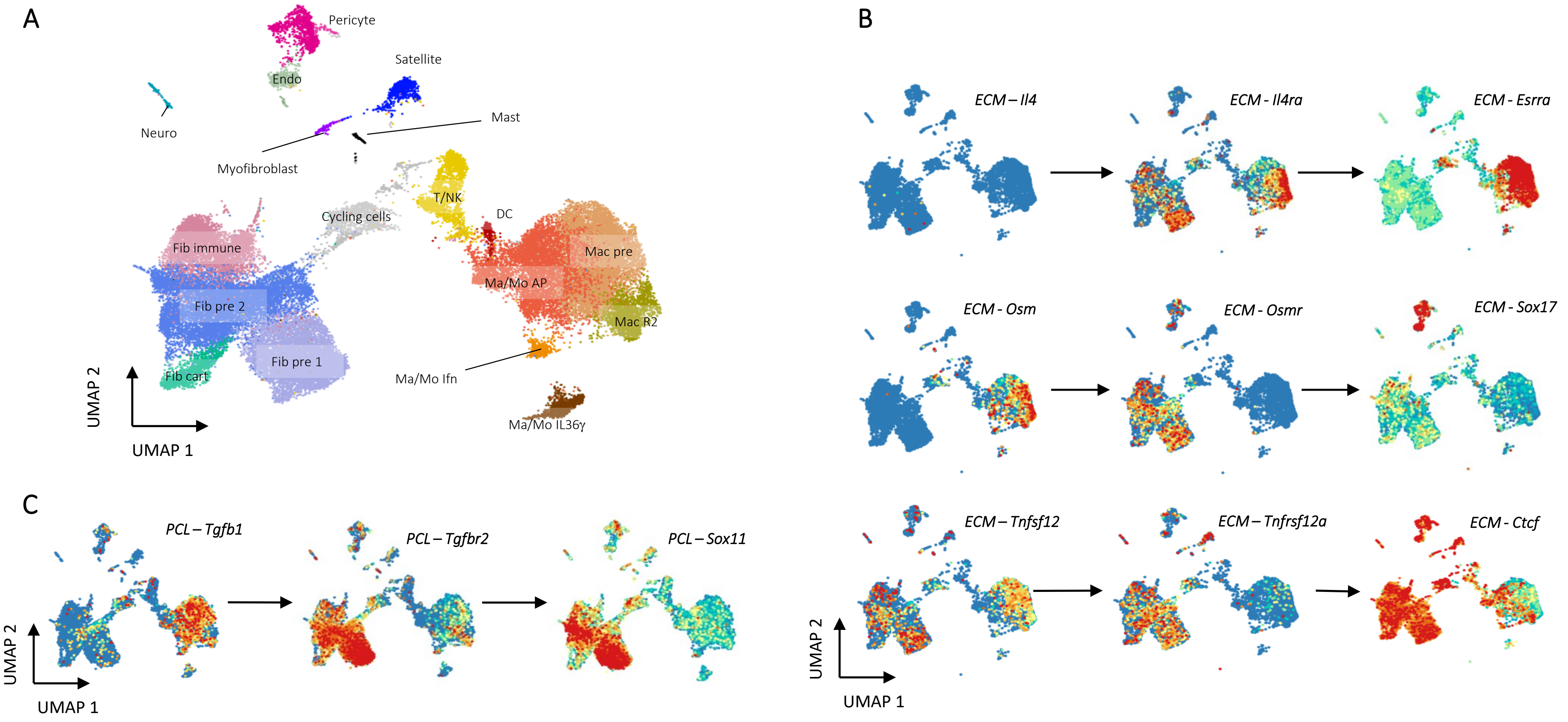
A
Pericyte
Satellite
Endo
Neuro
Mast
Myofibroblast
T/NK
Cycling cells
DC
Fib immune
Mac pre
Ma/Mo AP
Fib pre 2
Mac R2
Fib pre 1
Ma/Mo Ifn
Fib cart
UMAP 2
UMAP 1
Ma/Mo IL36γ
B
ECM – Il4
ECM - Il4ra
ECM - Esrra
ECM - Osm
ECM - Osmr
ECM - Sox17
ECM – Tnfsf12
ECM – Tnfrsf12a
ECM - Ctcf
UMAP 2
UMAP 1
C
PCL – Tgfb1
PCL – Tgfbr2
PCL – Sox11
UMAP 2
UMAP 1

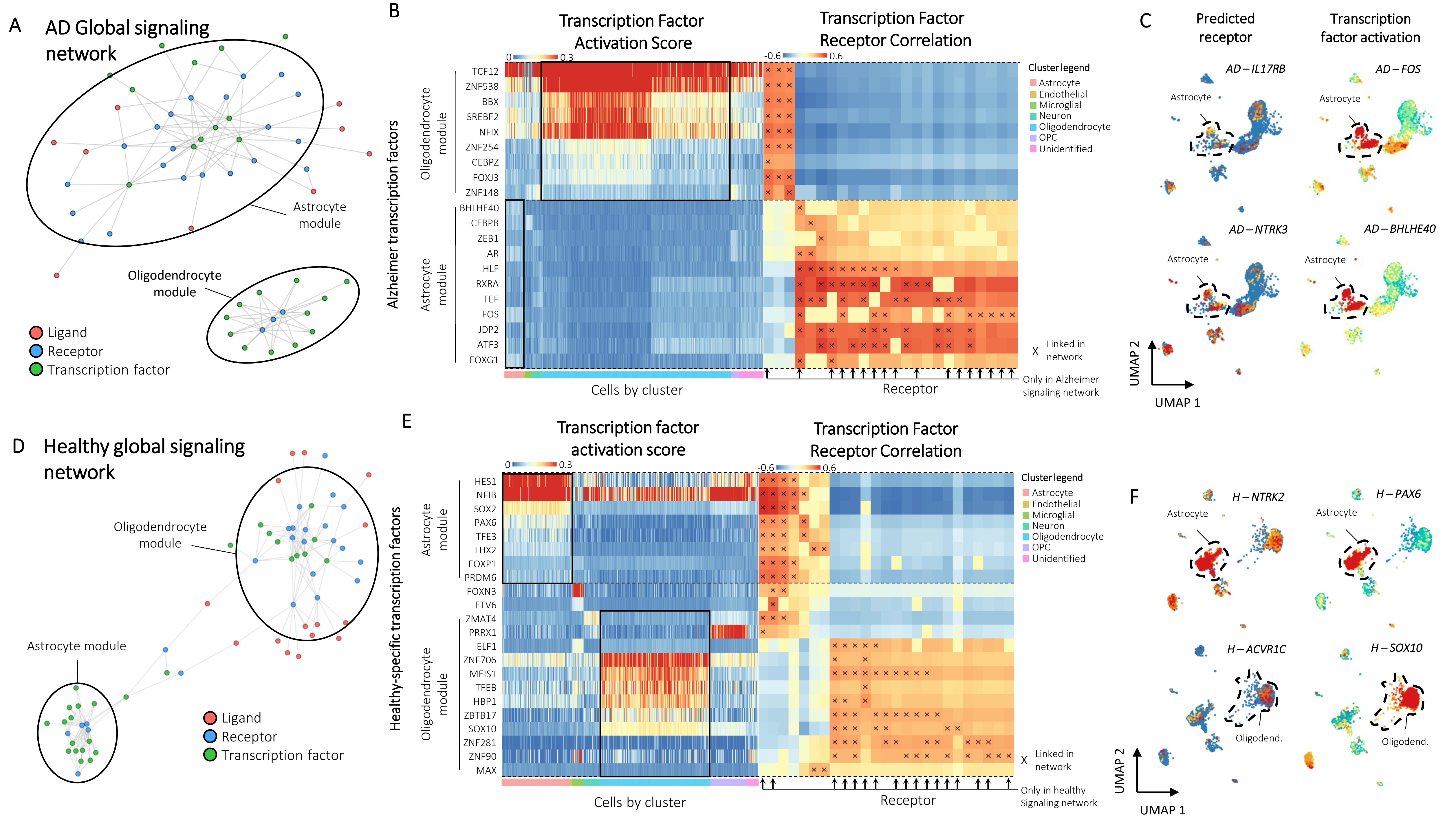

A
AD Global signaling network
Astrocyte module
Oligodendrocyte module
Ligand
Receptor
Transcription factor
B
Transcription Factor Activation Score
Transcription Factor Receptor Correlation
Alzheimer transcription factors
Oligodendrocyte module
Astrocyte module
TCF12
ZNF538
BBX
SREBF2
NFIX
ZNF254
CEBPZ
FOXJ3
ZNF148
BHLHE40
CEBPB
ZEB1
AR
HLF
RXRA
TEF
FOS
JDP2
ATF3
FOXG1
Cluster legend
Astrocyte
Endothelial
Microglial
Neuron
Oligodendrocyte
OPC
Unidentified
X Linked in network
Only in Alzheimer signaling network
Cells by cluster
Receptor
C
Predicted receptor
Transcription factor activation
AD – IL17RB
AD – FOS
AD – NTRK3
AD – BHLHE40
Astrocyte
UMAP 1
UMAP 2
D
Healthy global signaling network
Oligodendrocyte module
Astrocyte module
Ligand
Receptor
Transcription factor
E
Transcription factor activation score
Transcription Factor Receptor Correlation
Healthy-specific transcription factors
Astrocyte module
Oligodendrocyte module
HES1
NFIB
SOX2
PAX6
TFE3
LHX2
FOXP1
PRDM6
FOXN3
ETV6
ZMAT4
PRRX1
ELF1
ZNF706
MEIS1
TFEB
HBP1
ZBTB17
SOX10
ZNF281
ZNF90
MAX
Cluster legend
Astrocyte
Endothelial
Microglial
Neuron
Oligodendrocyte
OPC
Unidentified
X Linked in network
Only in healthy Signaling network
Cells by cluster
Receptor
F
H – NTRK2
H – PAX6
H – ACVR1C
H – SOX10
Astrocyte
Oligodend.
UMAP 1
UMAP 2
