## Supplementary figures and images for "Intercellular signaling dynamics from a single cell atlas of the biomaterials response"

### Supplementary File ECM signaling network.pdf

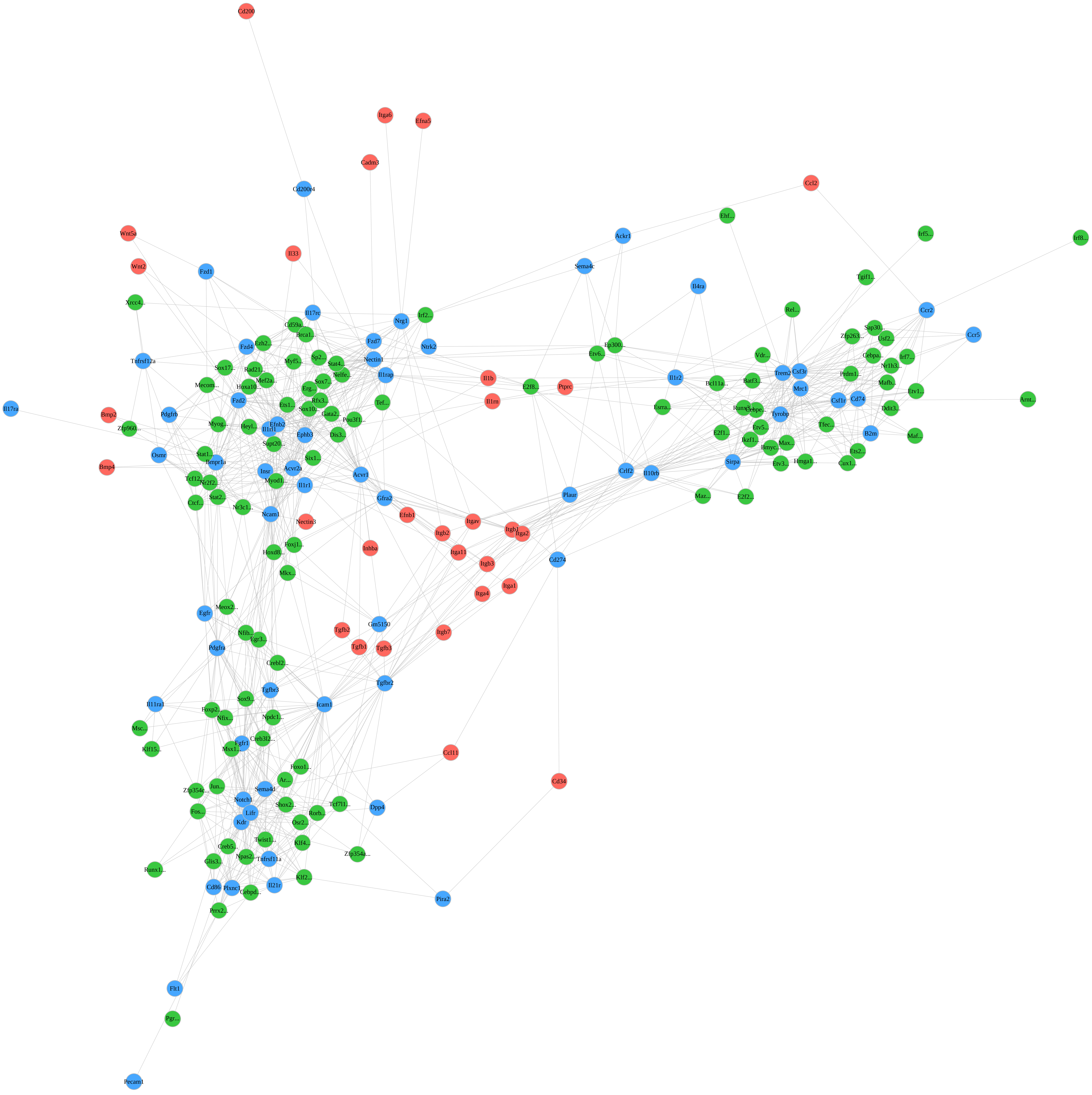

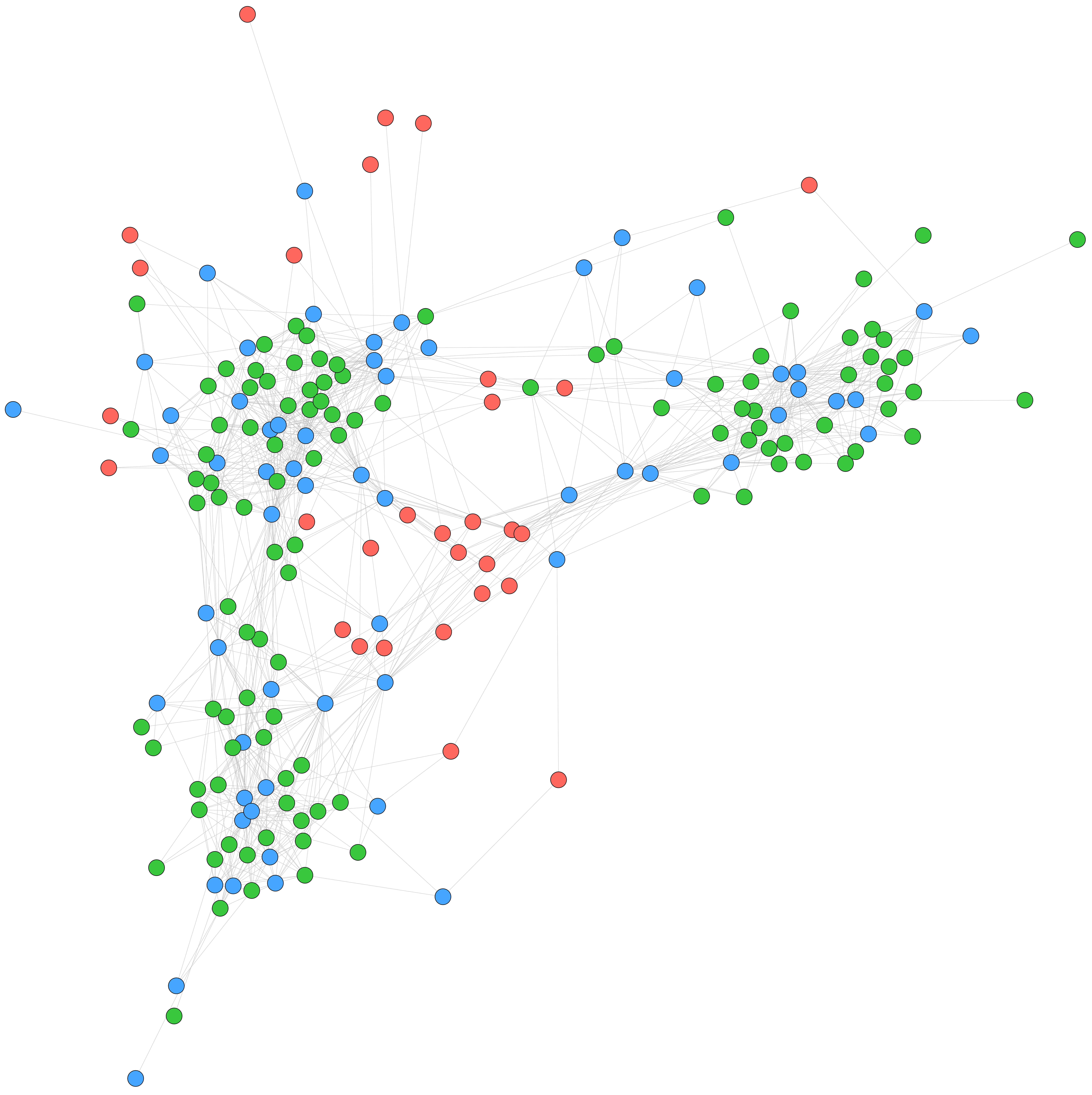

### Supplementary File PCL signaling network.pdf

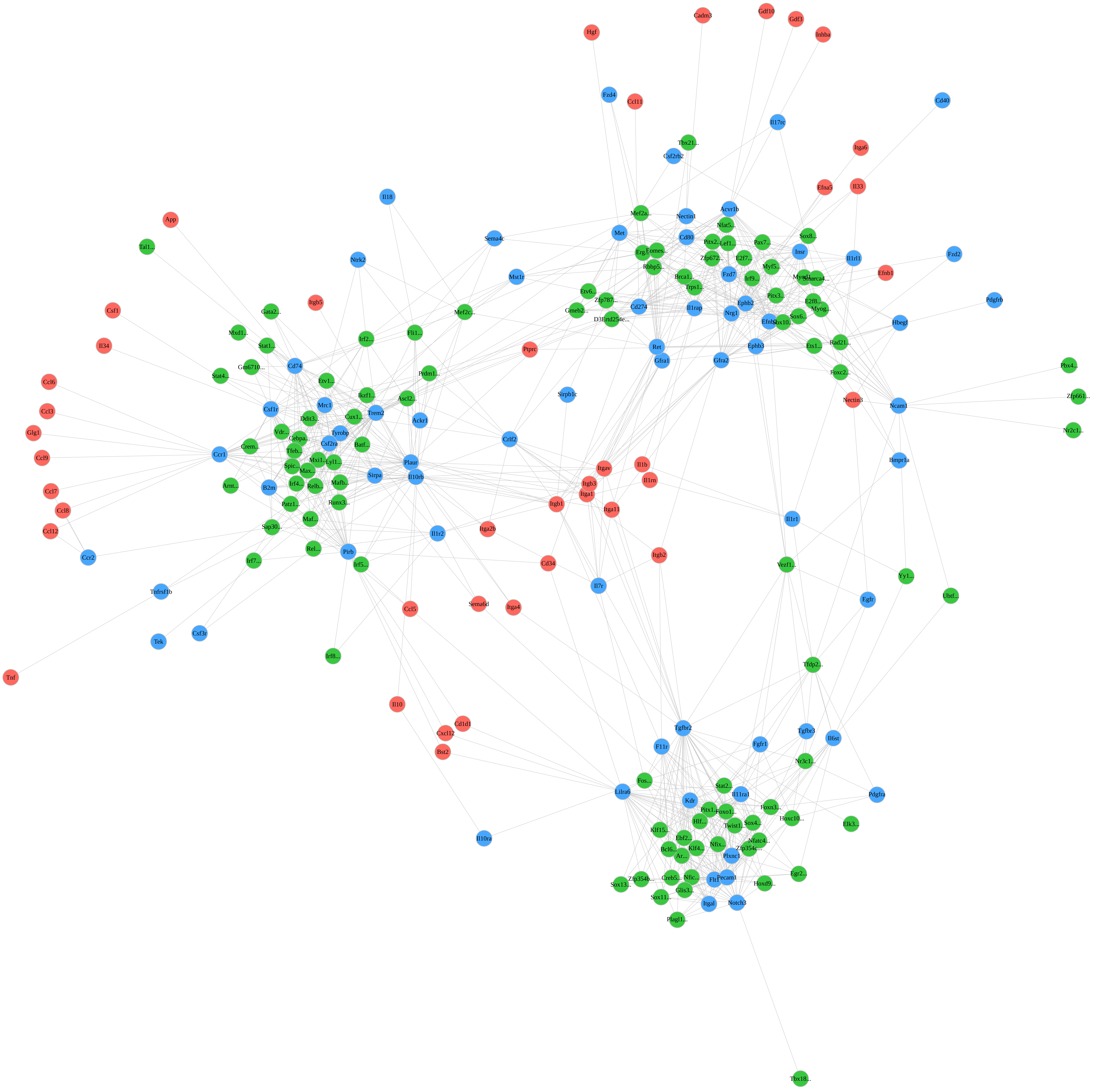

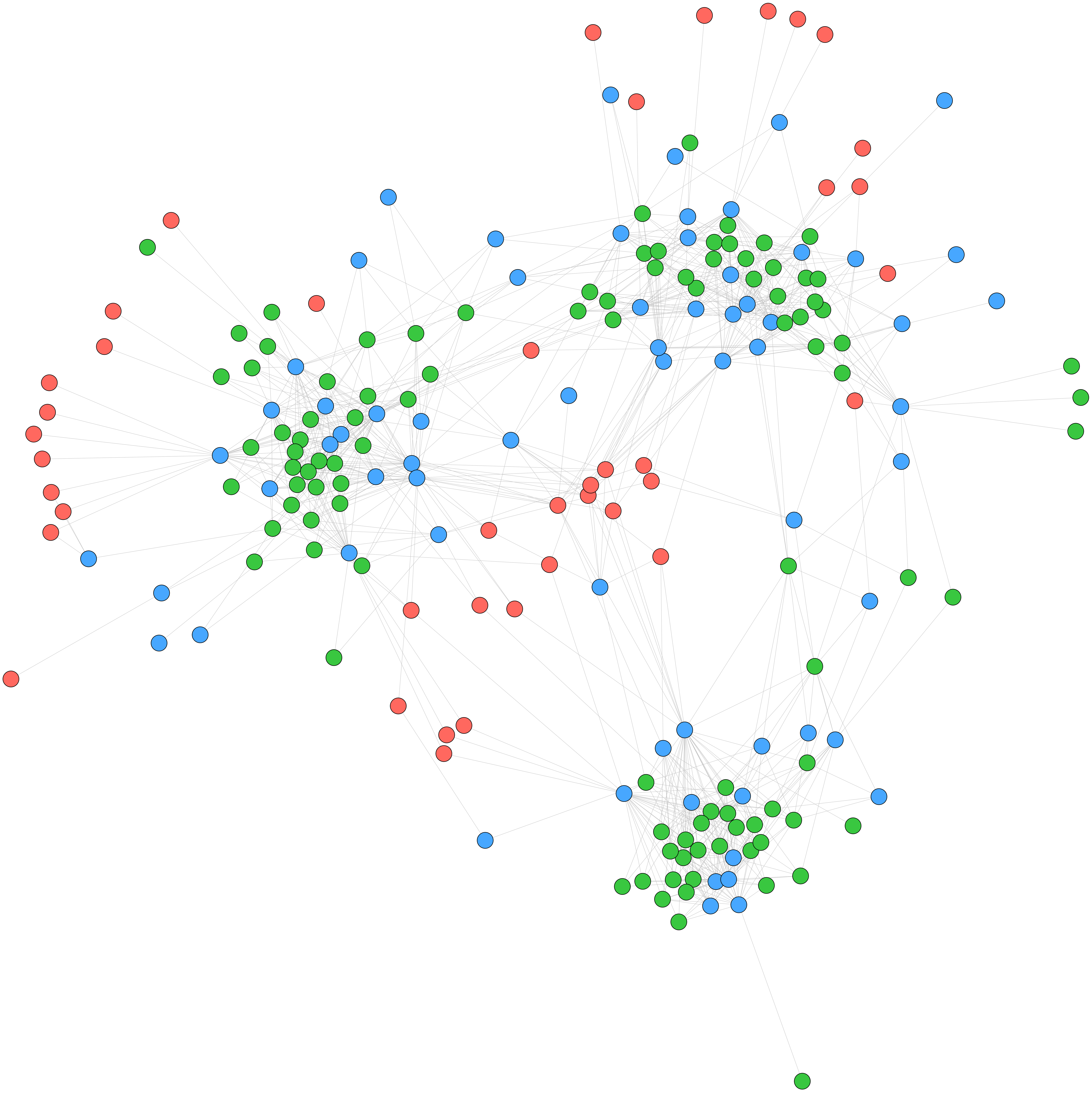
